# CRISPR/Cas9 knockout and editing of the major cysteine protease in *Entamoeba histolytica*

**DOI:** 10.64898/2026.06.08.731005

**Authors:** Wesley Huang, Rene L. Suleiman, Thomas Luder, Sylvia Jebiwott, Katherine S. Ralston

**Author notes:** Correspondence (K.S. Ralston).

## Abstract

Entamoeba histolytica is the cause of amoebiasis, a diarrheal infection that is a significant source of morbidity and mortality. Despite its importance to human health, the basic mechanism of disease is poorly understood. E. histolytica is dramatically understudied, even in comparison to other understudied parasites, due to challenging features of its genome and the relative paucity of genetic tools. A major limitation is the historical lack of gene knockouts or manipulation of the endogenous genome. While CRISPR/Cas9 has led to major advances in the tractability of numerous parasites, until recently, it was not applied in E. histolytica, except for the demonstration of editing of an episomal plasmid. In this study, we further developed CRISPR/Cas9 and demonstrated the successful targeting of the endogenous cysteine protease 5 (CP5) gene, a major virulence factor. Further, we demonstrated successful CRISPR/Cas9-based editing of this locus, which represents the very first successful editing of the endogenous genome. In the course of these approaches, we developed the viral skip peptide approach as well as a suite of endogenous promoters to drive tunable levels of gene expression. We also performed the most extensive empirical testing of nuclear localization signals in E. histolytica that has been done. Thus, this work demonstrates effective CRISPR/Cas9-based manipulation of the endogenous genome, together with significant advances to the overall genetic toolkit that extend beyond CRISPR/Cas9. These great improvements in the tractability of E. histolytica thus set the stage for future studies to better define its virulence.

## Introduction

Entamoeba histolytica is the cause of amoebiasis, a parasitic infection that is found predominantly in developing countries. Repeated diarrheal illness caused by E. histolytica has been associated with malnutrition and stunted growth in children [1, 2]. Clinically, amoebiasis can result in asymptomatic infection, diarrhea, bloody diarrhea with intestinal ulceration, or extra-intestinal abscesses that are fatal if not treated. It results in ∼50 million symptomatic infections per year, ∼15,500 deaths/year in children and 67,900 deaths/year in adults [3]. In children in a recent study, E. histolytica infection was associated with the highest risk of death [4]. Treatment options are limited to a single class of therapeutics, and there is no vaccine available [5]. Despite its importance to human health, the basic mechanism of disease is poorly understood. E. histolytica is dramatically understudied, even in comparison to other understudied parasites, due to challenging features of its genome and the relative paucity of genetic tools.

The E. histolytica genome is comprised of an estimated 38 linear chromosomes larger than 300 kb, and roughly 200 episomal DNA circles of about 25 kb that encode rRNA genes [6–8]. The genome is approximately 26.8 MB in size, with ∼75% A+T content and ∼8,745 genes [6]. There are numerous expanded gene families, comprising 56% of the proteome [9, 10]. No homologues have been identified for approximately one third of genes [7] and there are gene ontology (GO) terms for less than half of genes [9]. While ploidy is generally tetraploid, this varies at sub-chromosomal and chromosomal levels; hence, E. histolytica is considered to be aneuploid [6]. Genes involved in DNA repair and recombination are present in the genome, but experimental interrogation of these processes is limited. Genes involved in homologous recombination are present in the genome and the Rad51 homologue has been characterized in vitro [11]. Two studies have detected homologous recombination events either within plasmid episomes or the endogenous genome [12, 13]. Non-homologous end-joining (NHEJ) is not well characterized, and only the KU70 gene appears to be present, without an evident KU80 gene [14].

With regards to genetic tools, stable or transient transfection with circular plasmid DNA is a well-established approach [15–17]. Selectable markers on plasmids can be used to generate stable transfectants, in which plasmids are maintained as episomes [18]. Genes that are present on plasmid DNA are expressed exogenously, and this expression can be regulated through the use of tetracycline-inducible promoters or protein destabilization domains [19, 20]. Hence, exogeneous gene expression from plasmid DNA has been used as a tool to overexpress endogenous sequences or to express truncated or mutated gene sequences that are predicted to act as dominant negatives, e.g., [21]. These approaches have thus been used to interrogate gene function.

E. histolytica has an endogenous RNAi pathway that has also been exploited to interrogate gene function. Many strategies for RNAi knockdown have been applied experimentally. One such approach arose from the silencing of the amoebapore A gene through expression of its upstream sequences on a plasmid [22]. In this strain, known as “G3”, the expression of amoebapore A is knocked down, and additional genes can be knocked down by introducing new plasmids [22]. A caveat is that the expression of many other genes is altered [22] (see also dataset DS_1909b3c782 at amoebadb.org), making this approach fairly nonspecific. The more recently developed approach is the so-called “trigger” strategy [23]. In this approach, a plasmid is used to express a fragment of an endogenously silenced gene (the “trigger” sequence) fused to a fragment of the target gene [23]. Spreading of silencing leads to the production of small RNAs corresponding to the targeted gene, and stable knockdown of the endogenous copy of that gene [23]. This approach is very effective, particularly when clonal knockdown lines are obtained [24]. The trigger RNAi approach has also been adapted to create the first genome-wide E. histolytica RNAi library [25]. Despite the clear utility of RNAi, a major limitation is that there is no regulatable RNAi knockdown system in E. histolytica, making it less useful for the study of genes that are essential or that otherwise impact fitness.

The E. histolytica genetic toolkit has notably lacked methods to knockout endogenous genes. Also lacking are approaches to introduce sequences into the endogenous genome. Historically, the endogenous genome has not been successfully modified. In many other organisms, the application of the clustered regularly interspaced short palindromic repeat (CRISPR)/Cas9 system has been used for both targeted gene knockouts and genome editing. A related approach is CRISPR interference (CRISPRi), which enables targeted gene knockdown by impeding transcription with a catalytically inactive version Cas9 (dCas9) [26]. CRISPR/Cas9 and CRISPRi approaches have been successfully implemented to improve tractability of numerous parasites. CRISPR/Cas9 has been successfully applied in Cryptosporidium parvum [27], Plasmodium falciparum [28, 29], Plasmodium yoelii [30], Toxoplasma gondii [31, 32], Trypanosoma cruzi [33], Trypanosoma brucei [34, 35], Leishmania major [36], Leishmania donovani [37], Giardia lamblia [38] and Trichomonas vaginalis [39].

Until recently, CRISPR/Cas9 was not applied in E. histolytica. A key proof of concept for the application of CRISPR/Cas9 in E. histolytica was the demonstration of successful editing of an episomal plasmid [12]. In this work, a mutated luciferase gene on an episomal plasmid was successfully repaired by expressing Cas9 together with a guide RNA sequence and a repair template to restore the functional luciferase ORF [12]. These studies used regulated expression of Cas9 by making use of the destabilization domain from dihydrofolate reductase (ddDHFR) [19], as Cas9 expression proved to be toxic in E. histolytica. A key limitation of these studies was that an episomal sequence was edited, and there was no attempt to manipulate the endogenous genome.

Here we aimed to further develop CRISPR/Cas9 to target endogenous sequences, and to develop CRISPRi to complement available RNAi tools. A key advantage of CRISPRi is that the expression of dCas9 could be made regulatable, which would enable the first regulatable gene knockdown in E. histolytica. We demonstrate the successful editing of the endogenous cysteine protease 5 (CP5) gene using CRISPR/Cas9. This supports that DNA repair can occur via NHEJ. Further, when we provided a repair template, we demonstrated successful editing of the CP5 locus, supporting that DNA repair can also occur via homologous recombination. In the course of these approaches, we made significant advances to the overall genetic toolkit, including promoters for tunable expression, a viral skip peptide to link expression to selection, and empirical evaluation of nuclear localization signals (NLSs). These advances extend beyond the CRISPR/Cas9 approach. This work thus effectively develops CRISPR/Cas9, a powerful new approach for knockouts and editing of the endogenous genome, while also providing a wealth of broadly useful tools and advances that extend beyond CRISPR.

## Results

### The *E. histolytica* cysteine synthase promoter drives weak exogenous gene expression

To adapt CRISPRi for use in E. histolytica, we started by using the plasmid that was previously used to edit an episome in E. histolytica, pKT_Kan_ddCas9_NS-gRNA [12], which we will refer to as pCas9. This plasmid (Fig. S1a), contains the Streptococcus pyogenes Cas9 gene with an N-terminal Myc tag, Escherichia coli dihydrofolate reductase destabilization domain (ddDHFR) [19], and a putative E. histolytica nuclear localization signal (NLS3) [40]. The E. histolytica cysteine synthase (CS) promoter drives expression of Cas9. To adapt this plasmid for CRISPRi, site-directed mutagenesis was used to render the RuvC and HNH nuclease domains inactive, creating pdCas9 (Fig. S1b). For CRISPRi, high expression of dCas9 was desirable. Since destabilization domains can sometimes reduce gene expression levels, a version of this plasmid was created in which the ddDHFR destabilization domain was removed, creating pdCas9-constitutive (Fig. S1c).

E. histolytica trophozoites (“amoebae”) were stably transfected with pdCas9 or pdCas9-constitutive. As a control, amoebae were stably transfected with pKT-MG, an E. histolytica plasmid in which the CS promoter drives expression of Myc-tagged green fluorescent protein (GFP) (Fig. S1d). In E. histolytica, a common strategy to raise expression of exogeneous genes is to increase the amount of selective antibiotic. Thus, after obtaining stable transfectants, the level of selective antibiotic was increased prior to assaying for dCas9 expression. Immunofluorescence was performed using anti-Myc antibodies and samples were imaged by using imaging flow cytometry. The intensity of Myc staining in pdCas9 and pdCas9-constitutive transfectants was equivalent to the background level of Myc staining in wild-type control cells (Fig. S1e), thus, dCas9 expression was essentially undetectable. To further examine dCas9 expression, limiting dilution was used to obtain clonal lines. Among clonal lines, Myc staining remained equivalent to background (not shown). To determine if there was evidence for functional CRISPRi using these plasmids, two different guide sequences were used to target the cysteine protease 5 (CP5) gene for knockdown. Neither guide led to significant knockdown of CP5 expression in heterogeneous or clonal transfectants, as assessed by RT-qPCR (Fig. S1f). In these assays, we noted unusually variable expression of the housekeeping genes that are used in RT-qPCR analysis, which are typically highly stable.

Since Myc staining intensity in pKT-MG transfectants was only slightly higher than background (Fig. S1e), we reasoned that the CS promoter might drive weak exogenous gene expression. We also reasoned that although raising the amount of selective antibiotic is a common approach to raise exogenous gene expression in E. histolytica, the stress of increased antibiotic exposure might be causing variability in endogenous gene expression. In our hands, we have previously noted growth defects in stable transfectants in which G418 was increased. To further evaluate the impacts of increased G418, amoebae were stably transfected with pKT-MG (Fig. S2a). After obtaining stable transfectants, cells were grown with 6 μg/ml G418, the typical concentration used for maintenance of stable transfectants, or increased levels up to 50 μg/ml G418. In cells treated with 24 μg/ml or higher G418, gene expression became variable (Fig. S2b). In cells treated with 12 μg/ml or higher G418, cysteine protease activity was reduced (Fig. S2c). Thus, increased G418 is associated with pleiotropic deleterious effects and was not a desirable strategy to increase expression of dCas9.

### The P2A skip peptide and stronger endogenous promoters drive higher exogenous gene expression

Next, strategies were evaluated to increase exogenous gene expression without raising selective antibiotic. Viral 2A “skip” peptides such as P2A have been used in T. gondii to couple Cas9 expression and antibiotic resistance for improved stability and expression of Cas9 [41]. To test this approach in E. histolytica, Renilla Luciferase (RLUC) was used as a reporter gene. The pEhEx plasmid was modified such that P2A was used to couple the expression of the neomycin and RLUC ORFs, driven by either the E. histolytica CS or actin promoter (Fig. 1a), using plasmids referred to as pCS-RLUC-P2A and pActin-RLUC-P2A, respectively. As a control to assay RLUC expression driven by CS without P2A, pEhEx was modified such that RLUC was driven by the CS promoter and neomycin was driven by a different promoter (pCS-RLUC) (Fig. 1a). Stable transfectants were rarely recovered with pCS-RLUC-P2A, consistent with the relative weakness of the CS promoter. Stable transfectants were successfully recovered using pActin-RLUC-P2A, although it took notably longer than usual to recover stable transfectants. RLUC expression was approximately five-fold higher in pActin-RLUC-P2A transfectants than in pCS-RLUC transfectants (Fig. 1b), demonstrating that the viral skip peptide approach can be used effectively in E. histolytica.

**Fig. 1.**
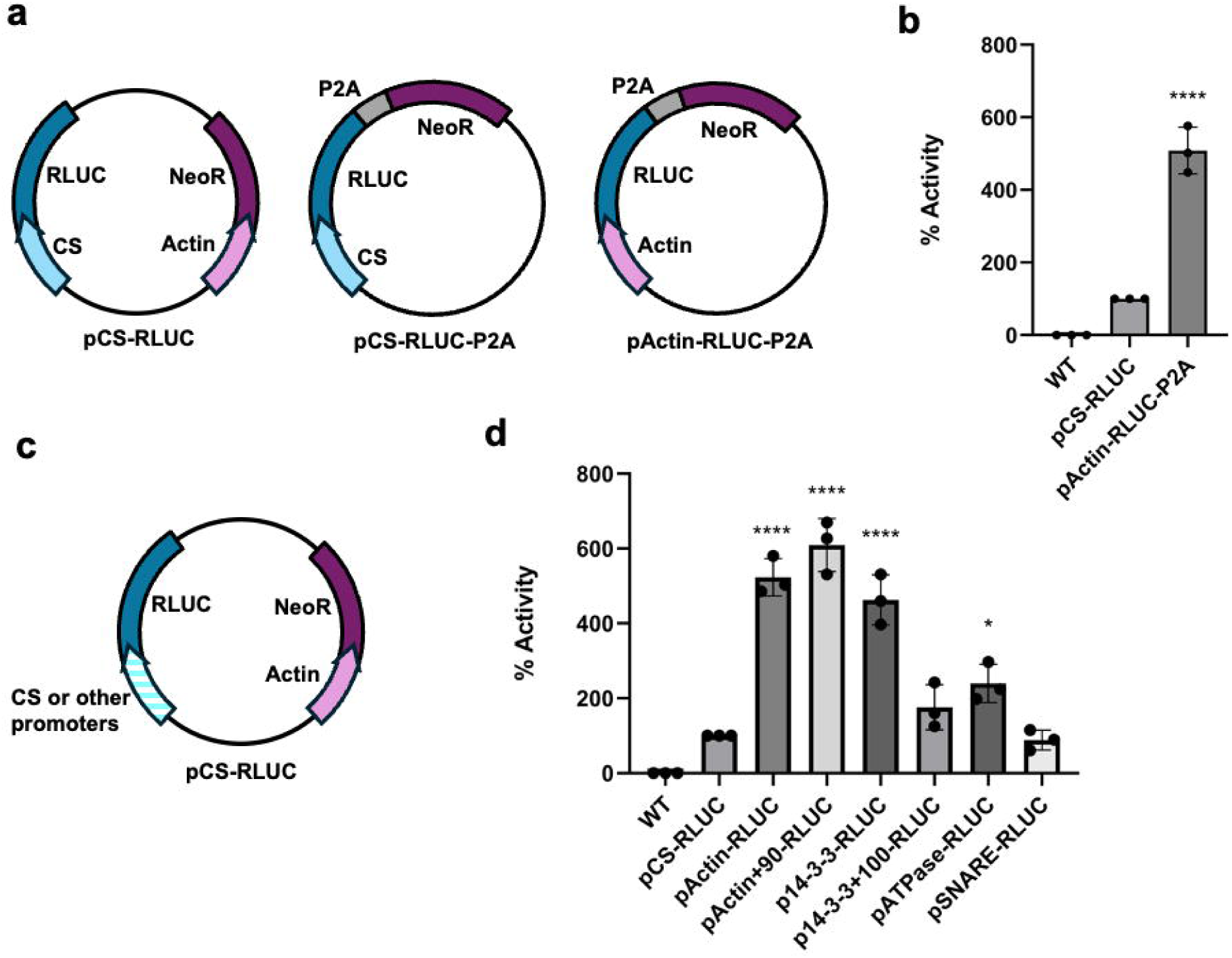
P2A and stronger endogenous promoters drive higher luciferase expression. **a**, pEhEx was modified such that RLUC was driven by the CS promoter (pCS-RLUC). pEhEx was also modified such that P2A was used to couple the expression of the neomycin and RLUC ORFs, driven by either the E. histolytica CS or actin promoter, creating pCS-RLUC-P2A and pActin-RLUC-P2A, respectively. **b**, Stable transfectants were assayed for luciferase activity using a Renilla Luciferase Assay kit. Control, wild-type parasites were set as 0%, and the activity of pCS-RLUC transfectants was set as 100%. **c**, pCS-RLUC was modified to replace the CS promoter with each candidate promoter sequence: actin, actin + 90, 14-3-3, 14-3-3 + 100, ATPase, or SNARE. **d**, Luciferase activity was assayed as in panel b. For panels b and d, n = 3, 3 independent experiments. 1-way ANOVA with Dunnett’s multiple comparisons test. *, p < .05, **, p < .01, ****, p < 0.001, ****, p < .0001.

In another approach, other endogenous E. histolytica promoters associated with highly expressed endogenous genes were tested. Gene expression data were used to identify candidate genes [42] and four were selected: Actin, 14-3-3, ATPase, and SNARE. For two of these genes, Actin and 14-3-3, downstream motifs within the ORF had been suggested to be necessary for promoter strength [42]. Thus, versions of the Actin and 14-3-3 promoters with and without the downstream motif (+90 and +100, respectively) were tested. pCS-RLUC was modified to replace the CS promoter with each candidate promoter sequence (Fig. 1c). Stable transfectants were readily obtained and Actin, Actin +90, 14-3-3, and ATPase promoters led to five- to six-fold higher RLUC expression than the CS promoter (Fig. 1d). The inclusion of the associated motifs (Actin +90 or 14-3-3 +100) was not required for increased RLUC expression compared to the absence of the motifs.

### Stronger endogenous promoters drive higher dCas9 expression and the ddDHFR destabilization domain enables regulatable expression

While the viral P2A skip peptide approach was effective in increasing exogeneous gene expression, owing to the slowness in recovering stable transfectants, we focused on using stronger promoters to drive dCas9 expression (Fig. 2a). The pdCas9 construct was modified such that the CS promoter was replaced with the actin, 14-3-3, ATPase or SNARE promoter. Western blotting analysis showed that all four promoters led to higher dCas9 expression than the CS promoter. Interestingly, while these constructs lacked the ddDHFR domain, there was no difficulty in recovering stable transfectants, and this high level of constitutive dCas9 expression appeared to be well tolerated. Next, the ddDHFR domain was added back to the pActin-dCas9 plasmid in order to enable regulatable dCas9 expression. Western blotting analysis demonstrated that dCas9 expression was induced by the addition of Trimethoprim (TMP) to stabilize ddDHFR (Fig. 2b).

**Fig. 2.**
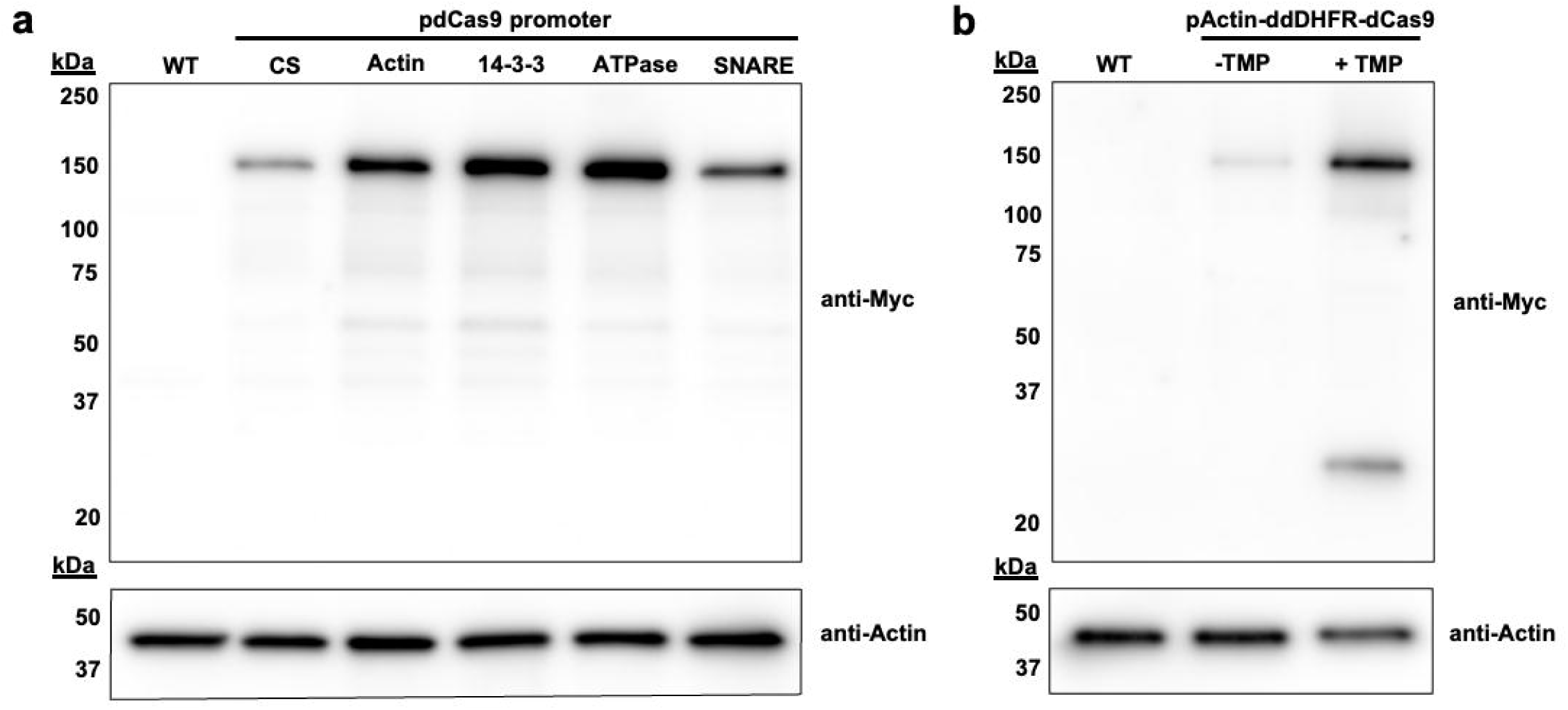
Stronger endogenous promoters drive higher dCas9 gene expression and the ddDHFR destabilization domain enables regulatable dCas9 expression. **a**, In the pdCas9 plasmid (see Fig. S1b), the CS promoter was replaced with actin, 14-3-3, ATPase, or SNARE promoters. Western blotting analysis of stable transfectants, or wild-type control parasites was used to assay for dCas9 expression. dCas9 was detected using an anti-Myc antibody (top panel), and an anti-actin antibody was used to control for loading (bottom panel). **b**, In the version of pdCas9 containing the actin promoter, the ddDHFR destabilization domain was re-introduced. As in panel a, Western blotting analysis was used to assay for dCas9 expression. Stable transfectants were induced with Trimethoprim (TMP) for 22-24 hours along with non TMP treated cells as control. Panel a is representative of two independent experiments and panel b is representative of two independent experiments.

### The 3x MIXed NLS localizes dCas9 to the nucleus and nuclear periphery

We next evaluated dCas9 localization. The endogenous E. histolytica NLS, NLS3, was used in a previous CRISPR/Cas9 study in E. histolytica, in which an episome was successfully edited [12]. In that study, NLS3 was at the N-terminus of Cas9. When we examined the localization of dCas9 with NLS3 at the N-terminus, immunofluorescence analysis demonstrated that dCas9 localized predominantly to the cytoplasm (Fig. S3). Thus, other NLSs were evaluated in order to localize dCas9 effectively to the nucleus.

Notably, NLSs in E. histolytica are poorly defined and only a few NLSs have been empirically evaluated. The DR rich tail of the E. histolytica Ago2-2 protein has a characterized non-classical NLS [43]. Thus, we added this DR-rich tail to the C-terminus of dCas9. We also tried adding another, longer, version of the DR-rich tail, “NLS/DR”, at the C-terminus of dCas9, but neither version of the DR-rich tail led to nuclear localization of dCas9 (Fig. 3). Next, we replaced the DR-rich tail with the SV40 NLS. Although the SV40 NLS has been used in E. histolytica [44], this NLS failed to localize dCas9 to the nucleus (Fig. 3). Subsequently, we replaced the N-terminal NLS3 with two fragments (position 17-178 and 314-333) from the endogenous RNA Pol II gene that are predicted to contain NLSs [45], but this did not lead to nuclear localization of dCas9 (Fig. 3).

**Fig. 3.**
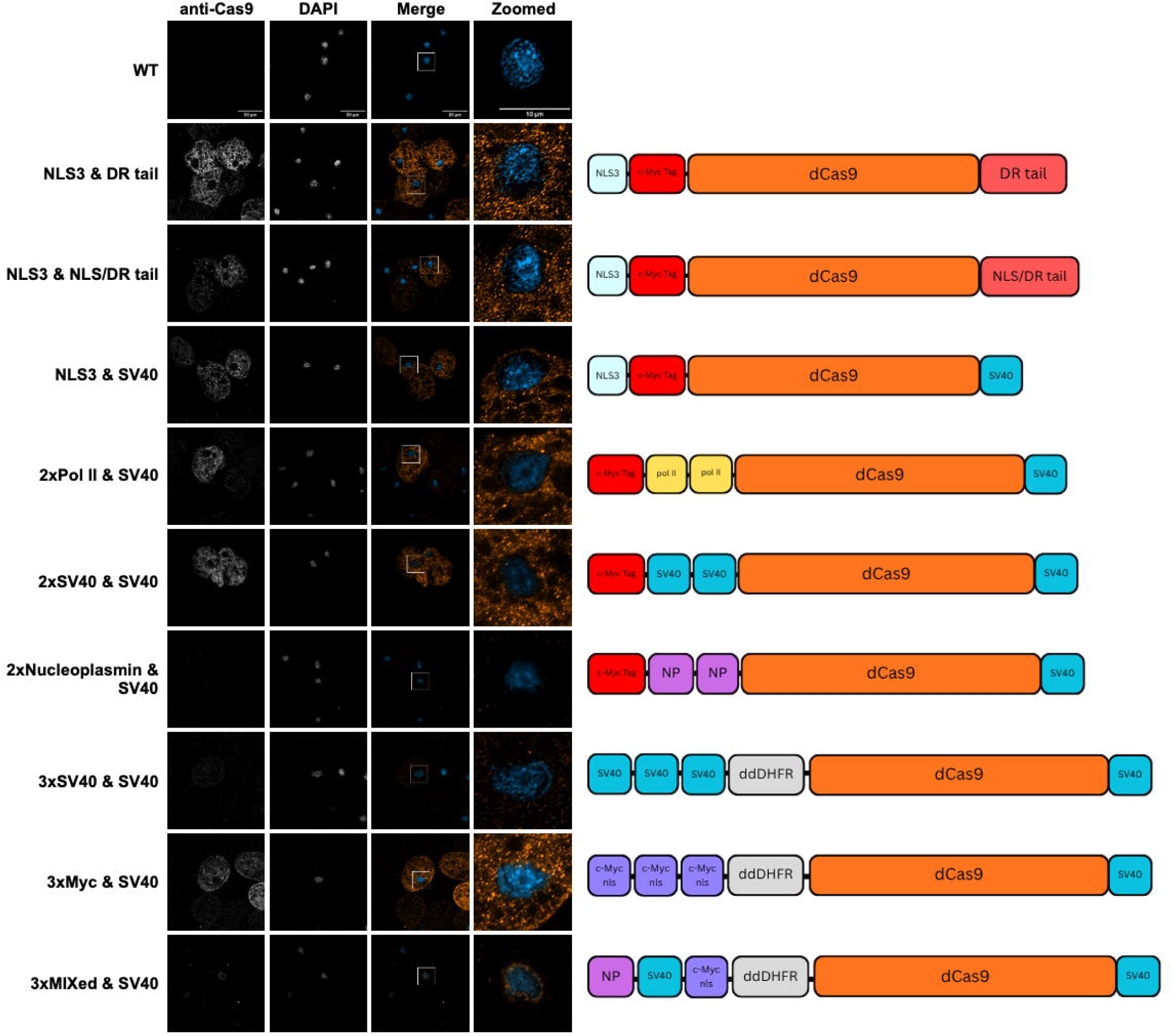
The 3xMixed NLS localizes dCas9 to the nucleus and nuclear periphery. To enable nuclear localization of dCas9, various NLSs were fused to the N- or C-terminus of dCas9, with or without the presence of an N-terminal ddDHFR destabilization domain. After obtaining stable transfectants, dCas9 was localized by using immunofluorescence with an anti-Cas9 antibody (orange), together with DAPI staining to visualize the nucleus (blue). Confocal images were captured on a Zeiss LSM 980 with Airyscan2. Wild-type amoebae were assayed as a control. The ddDHFR destabilization domain was re-introduced when designing the 3xNLS constructs; thus, these transfected amoebae were incubated with TMP overnight to induce dCas9 expression prior to immunofluorescence assays. Scale bar, 20 µm; zoomed images, 10 µm. Images are representative of 20-30 images collected for each stable transfectant line across two independent experiments.

In the course of this work, an in silico analysis of E. histolytica NLSs found that the most common monopartite and bipartite motifs were similar to the SV40 and Nucleoplasmin NLS, respectively [45], thus, we reasoned that these NLSs might be recognized by the putative endogenous importin proteins in E. histolytica. However, even with two tandem N-terminal Nucleoplasmin or SV40 NLSs, there was no evidence for nuclear localization, and the expression level of dCas9 with the Nucleoplasmin NLSs was poor (Fig. 3).

It is important to note that when we evaluated NLSs, the construct lacked the ddDHFR domain and dCas9 was thus constitutively expressed. It was possible that the lack of nuclear localization was due to potentially deleterious effects of constitutive dCas9 expression. Thus, the ddDHFR domain was reintroduced, and the Pol II, SV40, and Nucleoplasmin NLSs were evaluated again (Fig. S4). Nuclear localization of dCas9 was still not detected (Fig. S4a) even though the ddDHFR domain led to regulatable dCas9 expression (Fig. S4b).

It was possible that given the large size of dCas9, the NLSs that we had tested might be inaccessible to the putative endogenous importins. Alphafold was used to model dCas9 with candidate NLSs, together with putative E. histolytica importin alpha and beta sequences (Fig. S5). To make NLSs more accessible, new constructs were created with the NLS at the N-terminus, with a short linker between the NLS and ddDHFR, and a semi-rigid linker between ddDHFR and dCas9 (Fig. S5a). When this design was modeled using Alphafold, the NLSs appeared to be accessible to the putative importin alpha (Fig. S5b). Three different NLSs were tested: 3x SV40 NLS, 3x Myc NLS, and a 3x”MIXed” NLS (containing Nucleoplasmin, SV40, and Myc NLSs). The 3x SV40 NLS was associated with reduced dCas9 expression and did not lead to nuclear localization (Fig. 3). The 3x Myc NLS did not lead to nuclear localization (Fig. 3). By contrast, with the 3xMIXed NLS, dCas9 localized to the nucleus and nuclear periphery (Fig. 3). Finally, because the C-terminal SV40 NLS did not appear to aid in nuclear localization, it was removed. Localization of dCas9 to the nucleus was not affected by the removal of the SV40 NLS (Fig. S6). We will refer to this construct as p3xMIXed-dd-dCas9.

### CRISPR/Cas9 leads to effective knockout of the endogenous cysteine protease 5 gene

To test the effectiveness of CRISPRi, CP5 was targeted for knockdown because it is an established virulence factor and does not appear to be an essential gene [46]. While the E. histolytica genome contains at least 50 cysteine protease genes [10], CP5 is the most highly expressed in vitro [47]. As a positive control, CP5 was targeted for knockdown using the modern trigger RNAi system (Fig. S7). Clonal RNAi knockdown lines showed nearly undetectable CP5 gene expression by RT-qPCR analysis (Fig. S7a) and a concomitant significant decrease in total cysteine protease activity (Fig. S7b). Next, CP5 was targeted for CRISPRi knockdown using p3xMIXed-dd-dCas9 and three different gRNA sequences were tested, but RT-qPCR analysis did not reveal obvious evidence of CP5 knockdown (Fig. S8). This suggests that other aspects of CRISPRi may require further optimization.

At the same time as these experiments were carried out, we also simultaneously targeted CP5 for gene knockout by using active Cas9. Using p3xMIXed-dd-dCas9, site-directed mutagenesis was used to restore active Cas9, creating p3xMIXed-dd-Cas9. Like dCas9, Cas9 localized to the nucleus as expected (Fig. S6). Strikingly, when using active Cas9 to target CP5, RT-qPCR analysis demonstrated that CP5 levels in TMP-induced cells were dramatically reduced (Fig. 4a). When total cysteine protease activity was evaluated two days after TMP induction, cysteine protease activity was surprisingly increased (Fig. S9), suggesting that while this gene does not appear to be essential, there were compensatory changes in cysteine protease activity. When cysteine protease activity was evaluated two weeks after TMP induction, total cysteine protease activity was reduced (Fig. 4b).

**Fig. 4.**
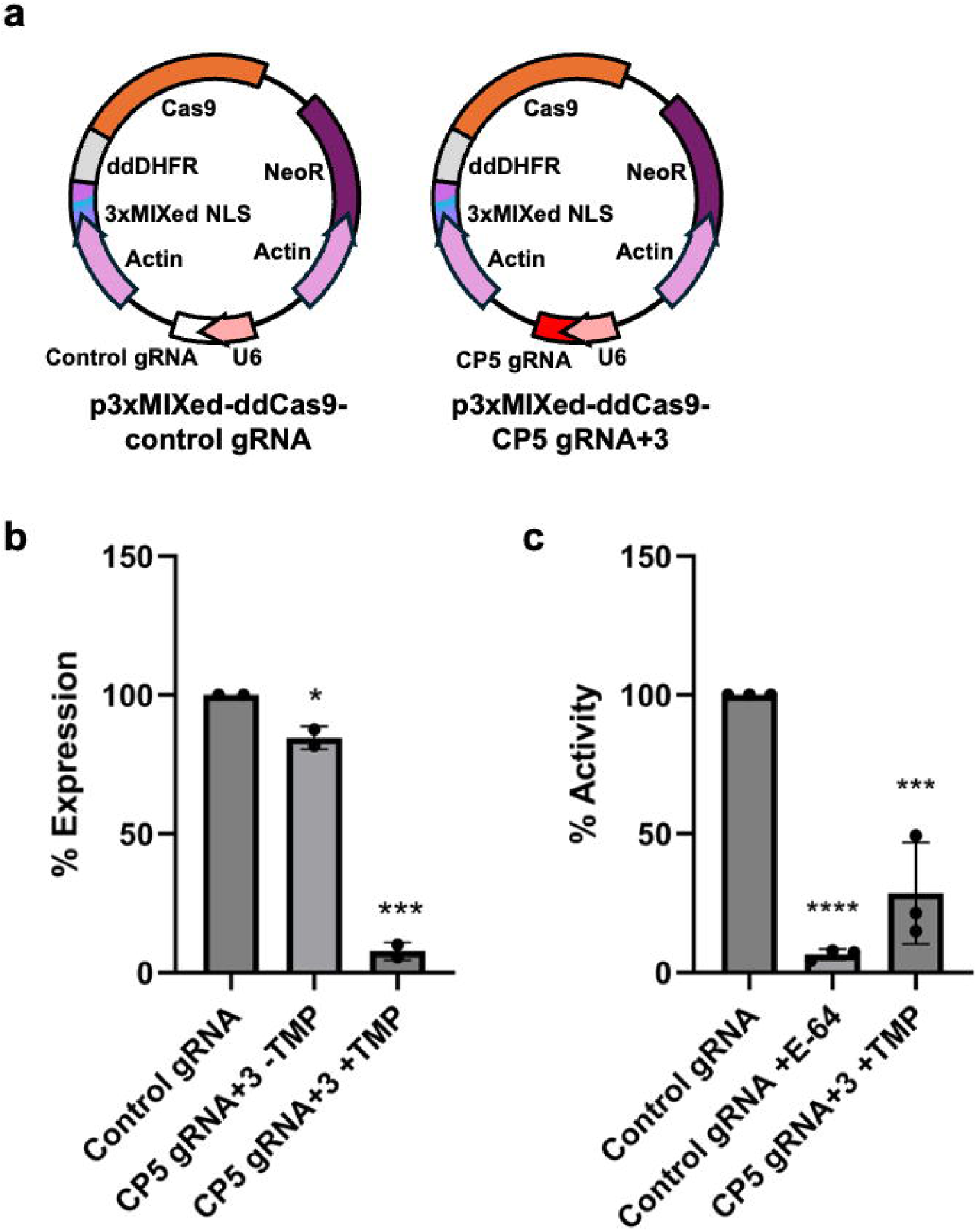
CRISPR/Cas9 leads to effective knockdown of CP5 expression. **a**, Amoebae were stably transfected with p3xMIXed-dd-Cas9, containing the actin promoter and active Cas9 with the 3XMixed NLS and ddDHFR destabilization domain. This plasmid also contained a nonspecific gRNA (Control gRNA) or gRNA targeting the CP5 ORF (CP5 gRNA+3). **b**, Stable transfectants were induced with TMP (or TMP was omitted, as a control) for two days to enable Cas9 expression prior to RNA extraction and RT-qPCR analysis of CP5 expression. CP5 expression was normalized to the RPL21 and VTP housekeeping genes. CP5 expression in control gRNA transfectants was set to 100%. N=2 separate RNA preps and two independent RT-qPCR assays. **c**, Stable transfectants were induced with TMP for two days to enable Cas9 expression, and then incubated in the absence of TMP for two weeks prior to assaying for total cysteine protease activity. Protease activity was assayed using Z-Arg-Arg-pNA substrate, and the activity of control gRNA transfectants was set to 100%. A separate control gRNA sample (Control gRNA) was incubated with the cysteine protease inhibitor E-64 to serve as an additional control. N=3, from 3 independent experiments. Panels b and c were analyzed by using 1-way ANOVA with Dunnett’s multiple comparisons test. *, p < .05, **, p < .01, ****, p < 0.001, ****, p < .0001.

### Editing of the CP5 locus using CRISPR/Cas9

To extend the CRISPR/Cas9 approach to editing of the endogenous genome, the CP5 locus was targeted. Amoebae were first stably transfected with p3xMIXed-dd-Cas9, containing either a nonspecific gRNA (vector control) or gRNA targeting the CP5 ORF (gCP5+3). After obtaining stable transfectants, a second plasmid, pCP5-homology, containing a region of CP5 (with the addition of a SpeI site) to serve as a repair template was introduced transiently. After adding TMP to induce Cas9 expression, genomic DNA was isolated and PCR was performed using primers specific to the endogenous CP5 locus (Fig. 5a). Digestion of PCR products with SpeI showed that the SpeI site was successfully introduced into the endogenous CP5 locus (Fig. 5b). The penetrance of the edited locus was high, with only a small proportion of the PCR product remaining uncut by SpeI. Thus, in addition to using CRISPR/Cas9 to target loci for the introduction of indels, CRISPR/Cas9 can be used to introduce targeted sequences into the endogenous genome.

**Fig 5.**
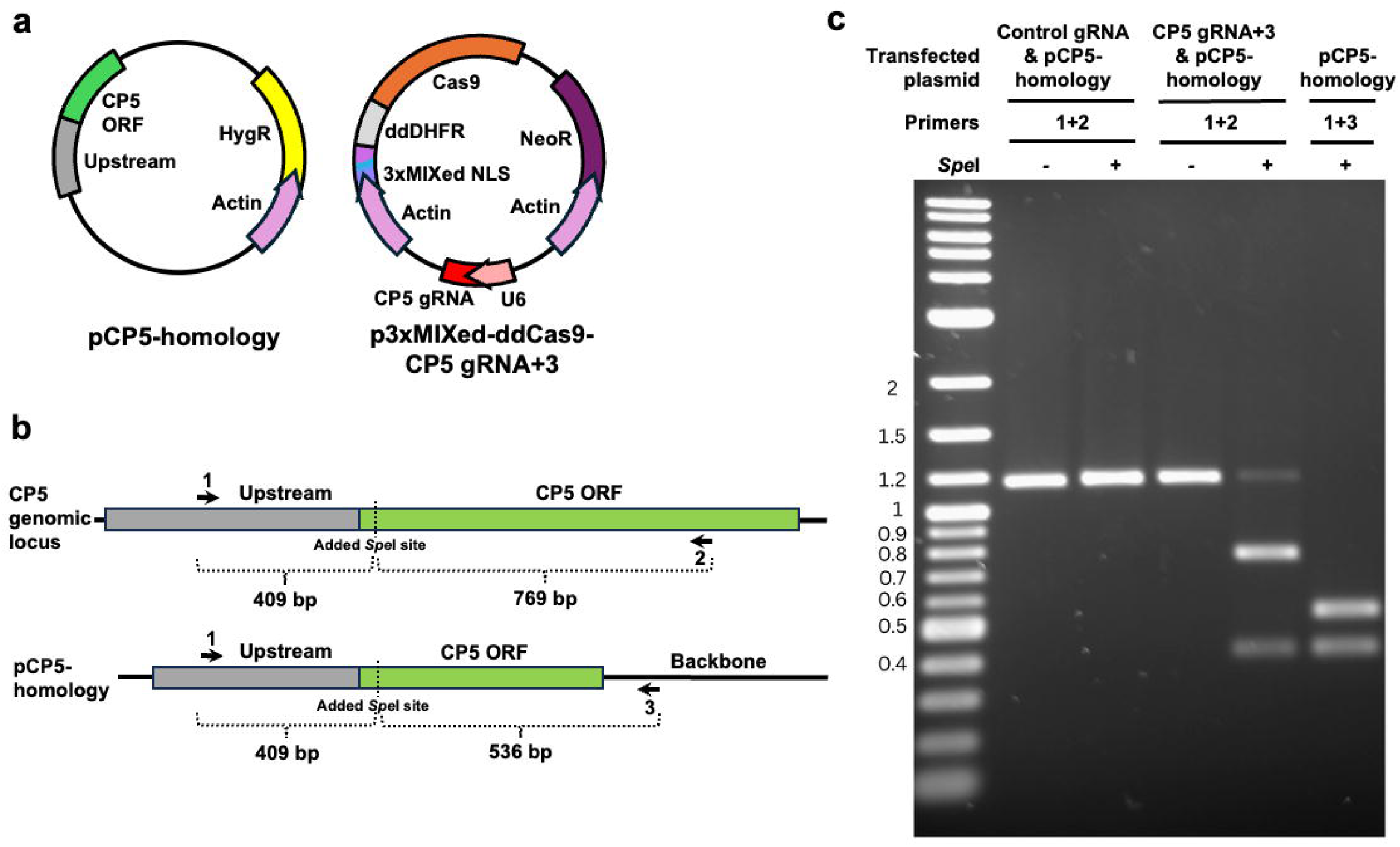
CRISPR/Cas9 can be used to introduce targeted sequences. **a**, Amoebae were stably transfected with p3xMIXed-dd-Cas9, containing the actin promoter and active Cas9 with the 3XMixed NLS and ddDHFR destabilization domain that co-expressed either a nonspecific gRNA (Control gRNA) or gRNA targeting the CP5 ORF (CP5 gRNA+3). Next, a second plasmid, pCP5-homology, was introduced transiently. This plasmid contains sequence homologous to the CP5 upstream and ORF sequences (with the addition of a SpeI site within the ORF) to serve as a repair template. **b**, Schematic diagram showing the positions of primers to amplify a fragment of the endogenous CP5 genomic locus (primers 1 + 2) or a fragment of the pCP5-homology plasmid (primers 1 + 3), and the position of the introduced SpeI restriction site is indicated. c, Genomic DNA was extracted from stable transfectants and PCR amplification was performed using the indicated primers. PCR products were incubated with or without SpeI. The CP5 genomic locus amplicon (primers 1 + 2) is 1,186 bp and digestion with SpeI produces 409 bp and 769 bp bands. The pCP5-homology amplicon (primers 1 + 3) is 945 bp and digestion with SpeI produces 409 bp and 536 bp bands.

## Discussion

These studies establish the first plasmid-based approach for CRISPR/Cas9 in E. histolytica. The endogenous genome was successfully manipulated to generate a knockout mutant, and successful editing of the endogenous genome was demonstrated for the first time. We further revealed the relatively weak expression level driven by the CS promoter, and developed the viral skip peptide approach as well as a suite of endogenous promoters to drive tunable levels of gene expression. By testing NLSs for the effective localization of dCas9/Cas9 to the nucleus, we also performed the most extensive experimental testing of NLSs in E. histolytica that has been done.

The CS promoter is commonly used to drive exogenous gene expression in a variety of E. histolytica plasmid constructs. Raising the concentration of selective antibiotic is a frequently used approach to increase exogenous gene expression, since this leads to an increase in plasmid copy number [18]. While G418 is often increased to levels of 50 μg/ml or higher, we found that even the small increase from 6 μg/ml to 12 μg/ml led to a marked decrease in cysteine protease activity, and an increase up to 24 μg/ml impacted gene expression. These findings suggest that increased G418 is an unfavorable method to increase exogeneous gene expression, owing to pleiotropic deleterious effects.

In this work, we developed several new approaches to modulate exogenous gene expression. By testing a set of endogenous promoters, we empirically uncovered promoters that drive varying levels of expression, expanding the molecular toolkit for exogenous gene expression. Although downstream motifs with the ORFs were thought to be necessary for promoter strength for two of these promoters, we did not find that these motifs were necessary for promoter strength. Beyond CRISPR/Cas9 approaches, these promoters are useful tools for the E. histolytica community, where each could be used to drive different desired levels of gene expression. Additionally, the viral P2A skip peptide is a new tool that can be applied for more stable exogenous gene expression, and to potentially increase exogenous gene expression. A limitation is that stable transfectants took longer to recover under selection, but this could possibly be remedied by using a stronger promoter, since the P2A approach could be combined with the use of stronger promoters.

Improving the regulation of exogenous gene expression in E. histolytica remains an ongoing goal. Destabilization domains including the ddDHFR destabilization domain have previously been established in E. histolytica [19], and are known to be associated with somewhat leaky gene regulation [43], as we also observed (Fig. 2b). Destabilization domains can potentially be used in tandem with existing tetracycline inducible approaches [20] to improve the tightness of regulatable expression. Other approaches such as the DiCre/loxP regulon [48] could be applied in the future. While Cas9 expression in E. histolytica was previously characterized as toxic, we observed no detrimental effects of constitutive dCas9 expression, and similarly did not have any difficulty in recovering or maintaining stable transfectants with Cas9 regulated by the somewhat leaky ddDHFR destabilization domain.

Aside from tuning dCas9 expression levels, the successful localization of this protein to the nucleus was a major hurdle. NLSs in E. histolytica have been poorly characterized and few have been tested experimentally. While the endogenous NLS3 sequence was used in the CRISPR/Cas9 proof of concept study in which an episomal sequence was successfully edited [12], the localization of Cas9 was not examined in that work. We found that dCas9 did not localize to the nucleus when using NLS3 (Fig. S3). NLS3 is a putative NLS from EhNCABP166, an actin-binding protein that has three putative bipartite NLSs [40]. It was shown that adding NLS3 to a fragment of the cytoplasmic EhFLN gene led to nuclear localization [40], but this was not true for dCas9, possibly owing to the much larger size of this protein. The lack of nuclear localization could be another reason that we did not see any evidence of gene knockdown when we initially tested CRISPRi using NLS3 to localize dCas9 to the nucleus (Fig. S1f). Given the limited knowledge of the amoebic nuclear import pathway and paucity of experimentally validated NLSs, we tested a putative NLS from the Ago 2-2 protein. The DR rich tail of the Ago2-2 protein is required for the nuclear localization of this protein and appeared to be sufficient to drive another DR tail-less Ago protein to the nucleus [43]. However, in the case of dCas9, the DR rich tail did not enable nuclear localization.

A recent in silico analysis of E. histolytica NLSs revealed that the most common monopartite and bipartite motifs were similar to the SV40 and Nucleoplasmin NLSs, respectively [45]. Thus we tested the SV40 or Nucleoplasmin NLSs, but did not detect nuclear dCas9 when using either of these NLSs individually. Because dCas9/Cas9 is a large protein, we reasoned that it could be more challenging to localize to the nucleus than smaller proteins, and that NLSs may require intervening sequences in order to be sufficiently accessible to importin proteins. Modeling studies using Alphafold supported the improved accessibility of NLSs to putative importins, when linker sequences were added (Fig S4). For dCas9, this approach, together with multiple different NLSs (Nucleoplasmin, SV40, Myc) led to successful nuclear localization. This suggests that the nuclear import pathway is conserved and intact and that common NLSs can drive nuclear import of target protein, even a relatively large protein like dCas9/Cas9.

In this work, we targeted CP5 for genetic manipulation because it is an established virulence factor, and crucially, it does not appear to be an essential gene. An existing CP5 knockdown mutant was made using the G3 strain many years ago [46]. This mutant has been continuously maintained in many laboratories for many years, with no loss of knockdown, suggesting it is not an essential gene. Interestingly, after initially inducing Cas9 expression to target the CP5 locus, we detected higher cysteine protease expression, suggesting that there was selective pressure to maintain protease activity, even though CP5 does not appear to be an essential gene. Ultimately, given more time, the level of cysteine protease activity was markedly reduced in the CP5 CRISPR/Cas9 mutant line. Notably, the reduction in overall cysteine protease activity (Fig. 4b) was much more pronounced than CP5 trigger RNAi mutants, even when there was nearly undetectable gene expression in those mutants (Fig. S7). The successful recovery of CP5 mutants is consistent with a functional NHEJ pathway, even though the genome appears to lack KU80 [14] and NHEJ has not yet been studied in E. histolytica.

In addition to using CRISPR/Cas9 to knockout CP5, we also successfully edited this locus. By providing a repair template containing a region of CP5, with the addition of a SpeI site, this sequence was successfully introduced into the endogenous CP5 locus. This is an important finding because it is the first demonstration of successful editing of the endogenous genome. Editing of the CP5 locus was highly efficient, since even in a bulk analysis of heterogeneous transfectants, only a minority of this locus remained unedited. These findings further support the presence of homologous recombination in E. histolytica and provide the first demonstration of homologous recombination of exogeneous sequences into the endogenous genome.

Until recently, CRISPR/Cas9 had not been applied in E. histolytica. A key advance was the demonstration of successful editing of an episomal plasmid [12]. The efficiency of editing was low [12], possibly due to the weak expression level of Cas9 driven by the CS promoter, and/or the ineffective localization of Cas9 driven by NLS3. While our studies were in progress, another group reported the successful knockout of an endogenous locus, ehcp112, in E. histolytica by using purified Cas9 in complex with gRNA as ribonucleoprotein (RNP) complexes [49]. The ehcp112 gene was knocked down to approximately 50% of the wild-type level, and indels were detected by Sanger sequencing of a PCR amplified gene fragment [49]. The penetrance of indels using this approach was not quantitatively examined. Immunofluorescence studies to localize Cas9 showed substantial cytoplasmic staining. The NLS that was used was not defined, thus there may be room for further improvements by modifying the NLS. The introduction of Cas9 RNPs has been an effective strategy in other organisms, and with further optimization, would complement the plasmid-based approaches defined in our studies. The RNP approach could potentially be applied relatively quickly, but the plasmid-based approach using stable transfectants allows for highly penetrant gene editing in a less heterogeneous population than the RNP strategy. Additionally, the plasmid-based approach allows for regulatable expression of Cas9, allows for gene editing by using a homologous repair template, and can be further optimized to enable CRISPRi in the future.

Our studies demonstrate successful gene knockout and gene editing of the endogenous genome using CRISPR/Cas9. This dramatically improves the tractability of E. histolytica. In addition, we made significant advances to the overall genetic toolkit that extend beyond the CRISPR/Cas9 approach, by developing new strategies for tunable exogenous gene expression and by empirically testing a variety of candidate NLSs. This work thus demonstrates the successful application of CRISPR knockouts and editing of the endogenous genome, while also providing a wealth of broadly useful tools and advances that extend beyond CRISPR. Together, we have vastly improved the ability to manipulate E. histolytica and set the stage for studies that will address the many open questions regarding its virulence.

## Materials and Methods

### Cell culture

The HM-1:IMSS E. histolytica strain (ATCC 30459) was cultured as described [24] in TYI-S-33 medium supplemented with 15% heat-inactivated adult bovine serum (Summa Life Sciences), 80 U/ml penicillin and 80 µg/ml streptomycin (Gibco), and Diamond Vitamin Tween 80 solution. The Diamond Vitamin Tween 80 solution was prepared according to Diamond et al., 1978 [50]. Cells were maintained in T25 tissue culture flasks (Corning) or glass culture tubes, and harvested for experiments at 80-90% confluence.

### DNA constructs

All plasmids generated for these studies were verified by restriction digest analysis and Sanger sequencing (Genewiz). Primers used for the these studies are found in Table S1.

We refer to pKT_Kan_ddCas9_NS-gRNA (Fig. S1a) [12] as pCas9. To inactivate Cas9, the QuikChange II XL Site-Directed Mutagenesis Kit (Agilent Technologies) was used. Two sequential rounds of mutagenesis were used to create D10A and H840A point mutations. The resulting plasmid is referred to as pdCas9.

To remove the ddDHFR domain in pdCas9, restriction digestion with AflII and AatII was used. After restriction digestion, gel extraction was used to recover the pdCas9 vector backbone using the QIAquick gel extraction kit (Qiagen). The pdCas9 vector backbone was modified using NEB Quick Blunting kit (NEB) and ligated using T4 DNA Ligase (NEB). The resulting plasmid is referred to as pdCas9-constitutive.

To use RLUC as a reporter gene, the RLUC ORF was cloned into the pEhEx [51] plasmid backbone. The RLUC sequence was amplified from pcDNA3 RLUC POLIRES FLUC. pcDNA3 RLUC POLIRES FLUC was a gift from Nahum Sonenberg (Addgene plasmid # 45642; http://n2t.net/addgene:45642; RRID:Addgene_45642) [52]. This PCR product was inserted into the PCR-amplified vector backbone [51] using the Gibson Assembly Ultra Kit (Synthetic Genomics). The resulting plasmid is referred to as pCS-RLUC.

To enable the use of the P2A skip peptide, the NeoR cassette first removed from the pEhEx-RLUC plasmid. This plasmid was digested with SpeI and EcoRI. After restriction digestion, gel extraction was used to recover the vector backbone using the QIAquick gel extraction kit. The vector backbone was modified using NEB Quick Blunting kit and ligated using T4 DNA Ligase. To introduce P2A-Neo downstream of the RLUC ORF, nested PCR was used to amplify Neo. The 5’ primer overhang contained the P2A sequence. This PCR product was inserted into the PCR-amplified vector backbone [51] using Gibson assembly. The resulting plasmid is referred to as pCS-RLUC-P2A. Finally, to replace the CS promoter with an actin promoter, the actin promoter was PCR amplified from pCS-RLUC and was inserted into the PCR-amplified vector backbone using Gibson assembly. Subsequently, the actin 3’ UTR was PCR amplified from pCS-RLUC and was inserted into the PCR-amplified vector backbone using Gibson assembly. The resulting plasmid is referred to as pActin-RLUC-P2A.

To replace the CS promoter in pCS-RLUC with various endogenous promoters, the CS 3’ UTR was first replaced with the actin 3’ UTR (associated with EHI_107290) using Gibson assembly. Next, approximately 500 bp upstream of the ORF of each corresponding gene Actin (EHI_107290), 14-3-3 (EHI_006810), VSNARE (EHI_081370), V-ATPase (EHI_059840) was PCR amplified and was inserted into the PCR-amplified vector backbone using Gibson assembly. The resulting plasmids are referred to as pActin-RLUC, p14-3-3-RLUC, pVSNARE-RLUC, and pV-ATPase-RLUC. Finally, to replace the CS promoter and UTRs in pdCas9, the actin promoter and 3’ UTR sequences were PCR amplified and inserted into the PCR amplified vector using two rounds of Gibson assembly. The resulting plasmid is referred to as pActin-dCas9.

To modify the NLS in pActin-dCas9, the various 2x NLS, and 3x/6x NLS with varying linkers and with or without the ddDHFR domain, were synthesized by Twist Bioscience. The version of this plasmid with the 3x MIXed NLS and the ddDHFR domain reintroduced is referred to as p3xMIX-dd-dCas9. Using the pdCas9 backbone, two sequential rounds of mutagenesis were used to repair the D10A and H840A point mutations, restoring the active Cas9 sequence. This plasmid is referred to as p3xMIX-dd-Cas9.

To insert gRNAs into eitherp3xMIX-dd-dCas9 or p3xMIX-dd-Cas9, the plasmid backbone was digested with BsaI and column purified using the QIAquick PCR purification kit (Qiagen). Single stranded gRNA oligos containing overhangs complimentary to the vector backbone were synthesized by Genewiz. Oligos were phosphorylated using T4 PNK (NEB) at 37°C for 30 minutes and after heat inactivation of T4 PNK, oligos were annealed with a ramp rate of 0.1°C/s to 25°C. Annealed oligos and restriction digested backbone were ligated with the Quick Ligation Kit (NEB).

To target CP5 for RNAi knockdown, a 512 bp region of the CP5 (EHI_168240) cds (position 202-714) was PCR amplified from genomic DNA. The 512 bp CP5 fragment was generated via a nested PCR approach, by first amplifying a larger 708 bp fragment. This PCR product was inserted into a SmaI-digested pTrigger vector [23] using Gibson assembly.

### 3D protein modeling

Modeling of dCas9 was performed in Alphafold3 through the Alphafold server (Google Ai). Putative E. histolytica importin alpha and beta (EHI_179940 and EHI_036520, respectively) was submitted along with dCas9 to the Alphafold server for structure prediction and interaction. The complex structure of dCas9 interacting with putative importin proteins was rendered in iCn3D (NCBI).

### Transfection

Amoebae were transfected using the Attractene transfection reagent (Qiagen) method as described previously [24]. Briefly, 20 µg of plasmid DNA was used for each transfection, and a no plasmid condition was included in each experiment as a control. For plasmids containing the Neo selectable marker, transfected amoebae were initially selected at 2 µg/mL G418 (Thermo Fisher Scientific). During selection, approximately half of the culture media was removed and replaced with fresh media containing G418, every 2-3 days. The G418 concentration was slowly increased to 3 µg/mL and then to 6 µg/mL. Stably transfected amoebae were then maintained at 6 µg/mL G418. Plasmids containing the Hygro selectable marker were transiently transfected to serve as a repair template for HDR mediated editing in amoebae that were already stably transfected with [plasmid name]. Hygromycin selection was not performed.

### Luciferase assays

The Renilla Luciferase assay kit (Promega) was used according to the manufacturer’s instructions. Stably transfected amoebae from 80-90% confluent T25 flasks were centrifuged at 200 x g for 5 minutes, then washed with 10mL 1x PBS once. Cell pellets were lysed in 500 µL of Renilla Luciferase Assay lysis buffer (Promega), supplemented with SigmaFast protease inhibitor diluted to 1x (Millipore). Cells in the lysis buffer were subjected to two rounds of freeze thaw to ensure complete cell lysis followed by brief vortexing. Cell debris was then removed by centrifugation at 16,000 x g for 2 minutes and supernatant was transferred to new microcentrifuge tubes. To measure protein concentration, 10 µL of each sample was tested with the Qubit Broad Range protein assay kit (Thermo Fisher Scientific) and measured on a Qubit 4 fluorometer (Thermo Fisher Scientific). 50 µg of total protein was loaded per technical replicate, totaling 3 technical replicates per condition on a black-walled 96-well plate with a flat, clear bottom (Corning). RLUC substrate was added immediately prior to measurement on the SpectraMax iD5 plate reader (Molecular Devices). Luminescence values were normalized to pCS-RLUC. Three independent experiments were performed.

### Protein extraction and Western blotting

Stable transfectants or wild type control cells were grown to 80-90% confluency in T25 flasks, and total protein samples were extracted. For constructs containing the ddDHFR domain, stable transfectants were incubated with Trimethoprim (TMP, Millipore Sigma) at a final concentration of 10 µM for 22-24 hours prior to protein extraction. For protein extraction, cells were centrifuged at 200 x g for 5 minutes at 4°C, then washed four times with ice cold 1x PBS. Cells were lysed in 1 ml RIPA lysis buffer (150mM NaCl, 1% Nonidet P-40, 0.5% Sodium Deoxycholate, 0.1% SDS, 50mM Tris pH 7.4) (Millipore Sigma) containing SigmaFast protease inhibitor (Millipore Sigma), with gentle agitation on a rocker at 4°C for 30 min. Samples were centrifuged at 16,000 x g for 5 minutes at 4°C. The supernatant was transferred to new tubes and protein concentration was measured using the Qubit 4 Fluorometer (Thermo Fisher). Using the Protein Broad Assay kit (Thermo Fisher), 10 µl of sample was added to 160 µl Protein BR assay buffer followed by addition of 30 µl Protein BR reagent in Qubit assay tubes (Thermo Fisher). Each sample was then briefly vortexed and incubated at room temperature, away from light, for 10 minutes prior to measurement.

To prepare the lysate for western blot, 250 µl of Laemmli sample buffer (0.25M Tris pH 6.8, 6% SDS, 40% Sucrose, 0.04% Bromophenol Blue (Thermo Fisher), 20% freshly added 2-mercaptoethnaol) (Millipore Sigma)) was added. Samples were heated at 70°C for 10 min, and cooled to room temperature prior to centrifugation at 16,000 x g for 2 minutes. For each sample, 20 µg of total protein supernatant was loaded onto pre-cast 4-20% polyacrylamide gradient gels (Bio-Rad). Gels were transferred onto nitrocellulose membranes (Bio-Rad) and then blocked in 1x PBS-T (1x PBS containing 0.05% Tween-20 (Millipore Sigma)) with 5% nonfat skim milk. Blots were probed at 4°C overnight on a rocker with either mouse monoclonal anti-Myc (9E10) (Thermo Fisher #13-2500) at a 1:1,000 dilution or mouse monoclonal anti-actin clone C4 (Millipore Sigma #MAB1501R) at a 1:1,000 dilution. Blots were washed four times with 1x PBS-T. Blots were then incubated for 1 hour at room temperature on a rocker, with goat anti-mouse IgG (H+L)-HRP conjugate secondary antibody (Bio-Rad #1721011), using a 1:25,000 dilution for anti-myc blots or a 1:5,000 dilution for anti-actin blots. Blots were washed four times with 1xPBS-T followed by one wash with 1x PBS. For detection, the clarity max ECL substrate (Bio-Rad) was used for anti-myc blots for higher sensitivity. Pierce ECL HRP substrate (Thermo Scientific) was used for anti-actin blots. Blots were imaged on an Amersham Imager 600 (GE Healthcare Life Sciences). Two independent experiments were performed using newly extracted protein samples each experiment.

### Immunofluorescence assays

1.2 cm diameter round coverslips (Electron Microscopy Sciences) were placed into 24-well plates (vendor), and coated with CellTak (Corning) according to the manufacturer’s instructions at 3.5 µg CellTak/cm^2^. Coverslips were washed 2 times with purified water and allowed to air dry. Stable transfectants or wild type control cells were grown to 80-90% confluency in T25 flasks. Cells were centrifuged at 200 x g for 5 minutes at 4°C and washed once in M199 medium (Gibco) supplemented with 5.7 mM cysteine, 25 mM HEPES, and 0.5% BSA (vendor). This medium is referred to as M199s. Cells were resuspended to 1×10^6^ cells/mL. and 100 µL was added to each coated coverslip in a 24-well plate. Plates were placed in an anaerobic GasPak (BD) and incubated at 35°C for 1 hour. Next, cells were fixed and permeabilized in 500 µL of ice cold 1:1 Methanol:Acetone for 10 minutes, and washed four times with 1x PBS. Coverslips were removed from the 24-well plates and placed inside humidity chambers. Coverslips were incubated with blocking solution (20% goat serum (vendor), 5% BSA (vendor) in 1x PBS) overnight at room temperature. Coverslips were washed four times in 1x PBS in quick succession and then washed four times in 1x PBS for 5 minutes per wash. Coverslips were probed with 50 µL of mouse monoclonal anti-Cas9 clone 7A9-3A3 (Thermo Fisher Scientific #MA1_201) at a 1:100 dilution for 3.5 hours at room temperature. Coverslips were washed four times in 1x PBS in quick succession and then washed four times in 1x PBS for 5 minutes per wash. Coverslips were incubated in 50 µL anti-mouse conjugated to Alexa Fluor 555 (Thermo Fisher #A-21422) at a 1:200 dilution for 3.5 hours in the dark. Coverslips were washed four times in 1x PBS in quick succession and then washed four times in 1x PBS for 5 minutes per wash. Coverslips were incubated in 1x PBS containing 1µg/ml 4′,6-diamidino-2-phenylindole (DAPI,Millipore Sigma) for 5 minutes and then washed once in 1x PBS. Washed coverslips were mounted onto microscope slides with 5 µL of Vectashield (Vector Laboratories) and sealed with nail polish. Samples were imaged on an LSM980 with Airyscan 2 (Zeiss) and images were processed on ImageJ. Two independent experiments were performed for each sample and approximately 10-15 images were collected across two coverslips in each independent experiment.)

### RNA extraction and RT-qPCR

Total RNA was extracted using the Direct-zol RNA purification kit (Zymo) following the manufacturer’s instructions. Stable transfectants or wild type control cells were grown to 80-90% confluency in T25 flasks. For each cell line, RNA extraction was performed on two separate days, per cell line to form biological replicate groups. Cells were centrifuged at 200 x g for 5 minutes at 4°C and the supernatant was removed. Cells were lysed in 300 µL of Trizol (Invitrogen), and then 300 µL of 100% ethanol (Millipore Sigma) was added. This mixture was transferred to the Direct-zol RNA purification kit binding column and washed with 400 µl RNA wash buffer. Bound samples were treated with DNase I and then washed with Direct-zol RNA Pre-Wash buffer. The samples were then washed with RNA wash buffer and finally eluted with 50 µL of RNase-free water.

For each sample, 2 µg of total RNA was resuspended in 20 µl RNase-free water and divided equally into two PCR tubes, one of which was used as a control lacking reverse transcriptase. For reverse transcription, SuperScript RT II (Invitrogen) was used according to the manufacturer’s instructions. For qPCR, cDNA samples were diluted by 1:1 or 1:10 with PCR grade water and added to aliquots of KAPA SYBR FAST qPCR Master Mix (Roche) containing separate primer sets. Two housekeeping genes, VTP (EHI_092140), and RPL21 (EHI_069110) were evaluated alongside the gene of interest. Technical duplicates of each sample were loaded onto a white LightCycler 96-well plate (Roche) and qPCR was carried out using a LightCycler480 II (Roche). Relative expression of the target gene was assessed using the 2^-ΔΔCt^ method. Two independent experiments were performed using RNA extracted on different days.

### Protease assays

Cysteine protease activity was measured as described in the literature [53, 54]. Stable transfectants expressing 3xMIX-dd-Cas9 and gCP5+3 were treated with TMP for 48 hours, and then cultured in the absence of TMP for two weeks prior to protease assays. Stable transfectants or wild type control cells were grown to 80-90% confluency in T25 flasks. Cells were centrifuged at 200 x g for 5 minutes at 4°C and washed twice with serum-free TYI-S-33. Washed cells were resuspended in serum-free TYI-S-33 to 5×10^4^cells/ml. 500 μl was transferred into a microcentrifuge tube and disrupted by four freeze-thaw cycles using an ethanol/CO_2_ bath. The resulting homogenate was centrifuged at 15,000 x g for 15 minutes at 4°C, and the supernatant was used for the cysteine protease activity assay.

Benzyloxycarbonyl-L-arginine-p-nitroanilide (Z-Arg-Arg-pNA • 2 HCl) was used as a substrate for measurement of protease activity (Bachem). The substrate was dissolved in 90% DMSO to a concentration of 10 mM. The assay buffer was prepared in purified water to the following specifications: 0.1M KHPO_4_/2mM EDTA/1mM DTT adjusted to pH 7.0 using potassium hydroxide. The assay was set up in a black-walled 96-well plate with a flat, clear bottom (Corning) in four replicates using 2 μl of substrate, 30 μl of cell lysate supernatant and 168 μl of assay buffer. As a control, lysate from wild type amoebae was incubated for 10 minutes at 37 °C with 50 μM E64 (N-[N-(L-3-trans-carboxyoxiran-2-carbonyl)-L-leucyl]-agmatine) (Sigma-Aldrich), a potent cysteine peptidase inhibitor. The rate of cleavage of the p-nitroaniline group was monitored over 30 minutes at 405 nm and 25°C using a SpectraMax iD5 plate reader (Molecular Devices). Relative enzyme activity was calculated using absorbance collected at the 20 minute timepoint. Three independent experiments were performed with samples extracted on three different days.

### Genomic DNA extraction, PCR, and *Spe*I digestion

Amoebic genomic DNA was extracted using the Zymo Quick DNA Miniprep Kit (Zymo Research) following the manufacturer’s instructions for cell monolayer samples. A Nanodrop 2000 (Thermo Fisher Scientific), was used to measure gDNA concentration. To prepare for PCR, the Q5 polymerase system was used (New England Biolabs) to make master mixes with each primer set for each gDNA sample. For each master mix sample, 90 ng of gDNA was mixed in and aliquoted into 8 PCR tubes calculated for 10 ng of gDNA per tube. After performing PCR, amplicons were then consolidated and purified using High Prep PCR beads (MagBio Genomics). In brief, an equal volume of beads was added to each consolidated PCR sample, mixed, and incubated at room temperature for 15 minutes prior to placing on a DynaMag-2 magnetic rack (Life Technologies). While on the magnetic rack, the supernatant was removed and the bead pellet was washed twice with 80% ethanol (Millipore Sigma). The bead pellet was air dried, resuspended in molecular grade water, and placed on the magnetic rack. Supernatant was transferred to clean tubes. 300 ng of purified amplicon was digested with 20 units of SpeI-HF in Cutsmart buffer (New England Biolabs). After 2 hours of incubation, an additional 20 units of SpeI-HF was added, and incubated for another 2 hours. Samples were visualized on a 0.8% agarose gel.

### Statistical analyses

All statistical analyses were performed using GraphPad Prism, and all data plots display means and standard deviation values. 1-way ANOVA with Dunnett’s multiple comparisons test was used for all statistical analyses. No statistical significance was indicated by a P of > 0.05. Statistical significance was indicated by *, P ≤ 0.05; **, P ≤ 0.01; ***, P ≤ 0.001; ****, P ≤ 0.0001.

## Acknowledgements

We thank Upi Singh (The University of Iowa) for the plasmid pKT_Kan_ddCas9_NS-gRNA that we refer to as pCas9 here. We thank Hannah Miller for creating pEhEx-RLUC and we thank Sara Tamayo-Correa for creating pTrigger-CP5. Imaging flow cytometry and microscopy were performed using shared instrumentation in the UC Davis MCB Light Microscopy Imaging Facility, of which the Zeiss 980 LSM was funded by NIH S10OD026702. We thank Thomas Wilkop for technical training and support in the use of these instruments. We thank the members of our laboratory, as well as Sean Collins and Priya Shah, for helpful discussions. This work was supported by NIH 1R01AI189918 and 1R01AI146914 awarded to K.S.R.. The funders had no role in study design, data collection and interpretation, or the decision to submit the work for publication.

## Author Contributions

W. H.: Methodology, Formal analysis, Investigation, Visualization, Writing – original draft, and Writing – review and editing. R.L.S.: Investigation. T.J.L.: Investigation. S.J.: Investigation. K.S.R.: Conceptualization, Funding acquisition, Project administration, Supervision, Visualization, Writing – original draft, and Writing – review and editing.

## Competing Interests

The authors declare no competing interests.

## Notes

### Competing Interest Statement

The authors have declared no competing interest.

