## Supplementary Information for "CRISPR/Cas9 knockout and editing of the major cysteine protease in *Entamoeba histolytica*"

**Supplementary Methods**

**Immunofluorescence assays and imaging flow cytometry**

To measure expression of Myc tagged proteins, cells were grown at 50 μg/ml G418 for 24 hours (see exposure of transfectants to increased G418, below). Cell lines transfected with constructs containing the ddDHFR domain were treated with 10 μM trimethoprim (TMP; Sigma-Aldrich) for 16-20 hours. Stable transfectants or wild type control cells were grown to 80-90% confluency in T25 flasks. Cells were centrifuged at 200 x *g* for 5 minutes at 4°C, washed twice in M199s, and resuspended to 7-9x10^6^ cells/ml in M199s. 500 μl was transferred to a microcentrifuge tube and fixed by incubation with 4% paraformaldehyde (Electron Microscopy Sciences) for 30 minutes at room temperature. After fixation, samples were washed once with 1x PBS and permeabilized in 1x PBS containing 0.2% Triton X-100 (Sigma-Aldrich) for 5 minutes. Samples were then washed three times in 1x PBS containing 0.1% Tween20 (1x PBST). Samples were incubated at 4°C on a rocker overnight in blocking solution (1x PBST containing 5% Bovine Serum Albumin (BSA; Gemini Bio-Products) and 20% Normal Goat Serum (Jackson ImmunoResearch Laboratories)). After blocking, samples were incubated with anti-Myc primary antibody (9E10; Abcam ab32) at a 1:50 dilution overnight at 4°C on a rocker. Samples were washed 4 times with 1x PBST and incubated at 4°C on a rocker for 3 hours with Cy5-conjugated AffiniPure Alpaca Anti-Mouse IgG (H+L) secondary antibody (1.5mg/ml; Jackson ImmunoResearch 615-175-214) at a 1:100 dilution. Samples were washed four times with 1x PBST and incubated with DAPI (Sigma) at a concentration of 40 µg/ml for 10 minutes. Samples were washed once in 1x PBS and resuspended in 1xPBS. Samples were assayed using a Luminex ImageStreamX MarkII imaging flow cytometer. 10,000 images were collected per sample.

**Exposure of stable transfectants to increased G418**

Stably transfected amoebae were routinely maintained at 6 µg/ml G418. To raise G418 concentrations prior to harvesting amoebae for experiments shown in Fig. S1 and S2, cells were passaged into new T25 flasks containing 12 µg/ml G418. Cells were incubated for one day followed by passaging into a new T25 flask containing µg/ml G418. This method was repeated to reach 50 µg/ml G418. Cells were incubated with 50 µg/ml G418 for one day prior to experiments.

**Immunofluorescence assays and confocal imaging**

Immunofluorescence assays were carried out as described in the Methods section, except that in addition to anti-Cas9 antibody, samples were probed with rabbit anti-H4 (Millipore #06-866) at a 1:500 dilution. For secondary antibodies, samples were incubated with both anti-mouse Alexa Fluor 555 (Thermo Fisher #A-21422) at a 1:200 dilution, and anti-rabbit Alexa Flour 488 (Jackson ImmunoResearch #111-545-003) at a 1:200 dilution.

**Supplementary Table Legends.**

**Table S1. Primers and oligos used in these studies.**

Sequences of primers and oligos that were used in these studies, for cloning of plasmid constructs, cloning of gRNA, site-directed mutagenesis of plasmid constructs, PCR, qPCR, and Sanger sequencing of plasmid constructs. The purpose of each primer is indicated, along with the primer name and sequence.

**Supplementary Figure Legends**

**Fig. S1. The CS promoter drives weak gene expression.**

**a,** We refer to pKT_Kan_ddCas9_NS-gRNA [12] as pCas9. This plasmid contains the *Streptococcus pyogenes* Cas9 gene with an N-terminal Myc tag, *E. coli* dihydrofolate reductase destabilization domain (ddDHFR) [19], and a putative *E. histolytica* nuclear localization signal (NLS3) [40]. The *E. histolytica* cysteine synthase (CS) promoter drives expression of Cas9. This plasmid also contains an endogenous U6 promoter together with a guide RNA corresponding to a sequence that is not present in the *E. histolytica* genome. **b,** The Cas9 gene in pCas9 was modified by using site-directed mutagenesis to inactivate Cas9, creating pdCas9. **c,** A version of pdCas9 was created that lacked the ddDHFR destabilization domain, pdCas9-consitutive. **d,** pKT-MG, an *E. histolytica* plasmid in which the CS promoter drives expression of Myc-tagged green fluorescent protein (GFP) was used as a control. **e,** Amoebae were stably transfected with pdCas9, pdCas9-constitutive or pKT-MG. After recovering stable transfectants at 6 μg/ml G418, the level of G418 was increased to 50 μg/ml. Immunofluorescence was performed using anti-Myc antibodies and samples were imaged by using imaging flow cytometry. Wild-type cells were analyzed as an additional control. One representative is shown, from two independent experiments. **f,** Amoebae were stably transfected with pdCas9-constitutive, containing a guide RNA corresponding to a sequence that is not present in the *E. histolytica* genome (Control gRNA) or a guide corresponding to CP5 +3 position (CP5 gRNA+3) or CP5 +385 position (CP5 gRNA+385). Clonal lines were obtained by limiting dilution. After recovering clonal lines, G418 was increased to 50 μg/ml and CP5 expression was evaluated by using RT-qPCR. CP5 expression was normalized to the RPL21 and VTP housekeeping genes. CP5 expression in control gRNA transfectants was set to 100% expression. N=2 separate RNA preps and two independent RT-qPCR assays.

**Fig. S2. High levels of G418 are deleterious.**

**a,** To evaluate the impacts of increased G418, amoebae were stably transfected with pKT-MG. **b,** After obtaining stable transfectants, cells were grown at 6 μg/ml G418, which is the typical concentration used for maintenance of stable transfectants. To increase G418, some cells were then grown with up to 50 μg/ml G418. RT-qPCR was used to assess CP5 expression, with expression was normalized to the RPL21 and VTP housekeeping genes. CP5 expression in WT cells was set to 100% expression. N=2 separate RNA preps and two independent RT-qPCR assays. **c,** Total cysteine protease activity was assayed using Z-Arg-Arg-pNA substrate , and the activity of wild-type parasites was set to 100%. A separate wild-type sample was incubated with the cysteine protease inhibitor E-64 to serve as an additional control. One representative is shown, from two independent experiments. 1-way ANOVA. ****, p < .0001, *, p < .05.

**Fig. S3. The existing NLS3 does not localize dCas9 to the nucleus.**

Amoebae were stably transfected with pdCas9-constitutive. The schematic shows the position of NLS3 and the c-Myc Tag in this construct. dCas9 was localized by using immunofluorescence with an anti-Cas9 antibody (orange), together with anti-histone H4 antibody (green) and DAPI staining (blue) to visualize the nucleus. Confocal images were captured on a Zeiss LSM 980 with Airyscan2. Wild-type amoebae were assayed as a control. Scale bar, 20 µm. Images are representative of 20-30 images of each cell line from two independent experiments.

**Fig. S4. Using the ddDHFR destabilization domain to regulate 2xNLS-dCas9 expression does not lead to nuclear dCas9 localization.**

Downstream of the c-Myc Tag, the ddDHFR destabilization domain was reintroduced and NLS3 was replaced with either two tandem NLSs from Pol II, SV40, or Nucleoplasmin (NP). At the C-terminus of dCas9, an SV40 NLS was added. **a,** Stably transfected amoebae were incubated with TMP overnight and dCas9 was localized by using immunofluorescence with an anti-Cas9 antibody (orange) and DAPI staining (blue) to visualize the nucleus. Confocal images were captured on a Zeiss LSM 980 with Airyscan2. Wild-type amoebae were assayed as a control. Scale bar, 20 µm; zoomed panels,10 µm. **b,** Amoebae stably transfected with the 2xSV40 NLS construct shown in panel a were incubated in the presence or absence of TMP overnight. Scale bar, 20 µm. Images are representative of 20 images of each cell line from two independent experiments.

**Fig. S5.Structural modeling of dCas9 with the ddDHFR destabilization domain and 3xMIXed NLS.**

dCas9 with an added 3xNLS and ddDHFR destabilization domain was modeled using the AlphaFold server. The predicted model was rendered in iCn3D. **a,** Rendered model showing dCas9 with the 3xMIXed NLS and ddDHFR domain. The semi-rigid linker (white) is located between N-terminus of dCas9 (magenta) and the ddDHFR domain (“dd”, green). Other versions of the 3xNLS were 3xSV40 and 3XMyc, which used the same spacers as shown here. **b**, dCas9 (magenta) with an added 3xMIXed NLS (turquoise) and ddDHFR domain (green) was modeled together with putative *E. histolytica* importin alpha (blue) and beta (brown).

**Fig. S6. The 3xMIXed NLS localizes dCas9 to the nucleus and nuclear periphery, and the C-terminal SV40 NLS is not required.**

Amoebae were stably transfected with p3xMIXed-dd-dCas9 or p3xMIXed-dd-Cas9. These constructs contain the N-terminal 3xMIXed NLS and lack the C-terminal SV40 NLS. After obtaining stable transfectants, amoebae were incubated with TMP overnight to induce dCas9/Cas9 expression prior to immunofluorescence assays. dCas9 and Cas9 were localized by using an anti-Cas9 antibody (orange), together with DAPI staining to visualize the nucleus (blue). Confocal images were captured on a Zeiss LSM 980 with Airyscan2. Scale bar, 20 µm. Images are representative of one independent experiment.

**Fig. S7. CP5 can be knocked down using the trigger RNAi system.**

A fragment of CP5 was cloned into the pTrigger construct to enable RNAi knockdown of CP5 as a control. Amoebae were stably transfected with pTrigger-CP5 (CP5 RNAi), or an empty pTrigger plasmid (Vector) as a control. After obtaining heterogeneous stable transfectants, clonal pTrigger-CP5 transfectants were obtained by limiting dilution. **a,** RT-qPCR was used to assess CP5 expression, with expression normalized to the RPL21 and VTP housekeeping genes. CP5 expression in control pTrigger (Vector) transfectants was set to 100% expression. Heterogeneous pTrigger-CP5 transfectants (Het.) and two different clonal lines (Clone 1 and Clone 2) were assessed. N=2 separate RNA preps per cell line and two independent RT-qPCR assays. **b,** Total cysteine protease activity was assayed using Z-Arg-Arg-pNA substrate, and the activity of pTrigger (Vector) transfectants was set to 100%. n=3 technical replicates, from 3 independent experiments.1-way ANOVA with Dunnett’s multiple comparisons test. *, p < .05, **, p < .01, ****, p < 0.001, ****, p < .0001.

**Fig. S8. No evidence for knockdown of CP5 expression using CRISPRi.**

Amoebae were stably transfected with p3xMIXed-dd-dCas9, containing a guide RNA corresponding to a sequence that is not present in the *E. histolytica* genome (Control gRNA) or a guide corresponding to CP5 +3, +183 or +256 positions (CP5 gRNA+3, +183 or +256). As a control, amoebae were stably transfected with pTrigger-CP5 (CP5 RNAi). With the exception of pTrigger-CP5 transfectants, transfected amoebae were treated with TMP for 2 days prior to RNA extraction. RT-qPCR was used to assess CP5 expression, with expression normalized to the RPL21 and VTP housekeeping genes. CP5 expression in control gRNA transfectants was set to 100% expression. N=2 separate RNA preps per cell line and two independent RT-qPCR assays. 1-way ANOVA with Dunnett’s multiple comparisons test. *, p < .05, **, p < .01, ****, p < 0.001, ****, p < .0001.

**Fig. S9. Initially higher CP activity immediately after Cas9 induction.**

Amoebae were stably transfected with p3xMIXed-dd-Cas9, containing a guide RNA corresponding to a sequence that is not present in the *E. histolytica* genome (Control gRNA) or a guide corresponding to CP5 +3 position (CP5 gRNA+3). Stable transfectants were treated with TMP for two days and then assayed for total cysteine protease activity. Protease activity was assayed using Z-Arg-Arg-pNA substrate, and the activity of vector control gRNA transfectants was set to 100%. A separate control gRNA sample was incubated with the cysteine protease inhibitor E-64 to serve as an additional control. N=3, from 3 independent experiments. 1-way ANOVA with Dunnett’s multiple comparisons test. *, p < .05, **, p < .01, ****, p < 0.001, ****, p < .0001.
