## Supplementary figures and images for "CRISPR/Cas9 knockout and editing of the major cysteine protease in *Entamoeba histolytica*"

### Figure S1

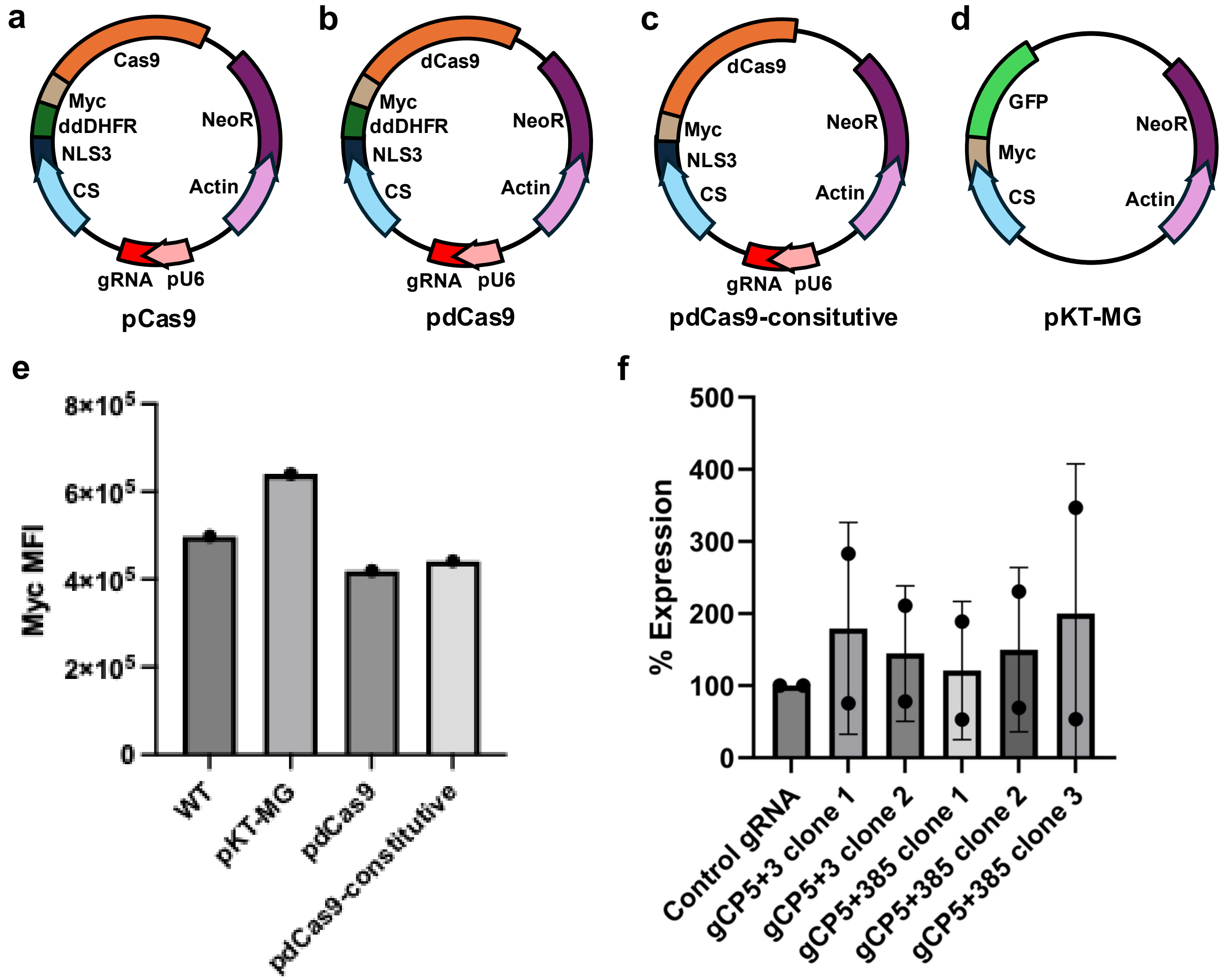

### Figure S2

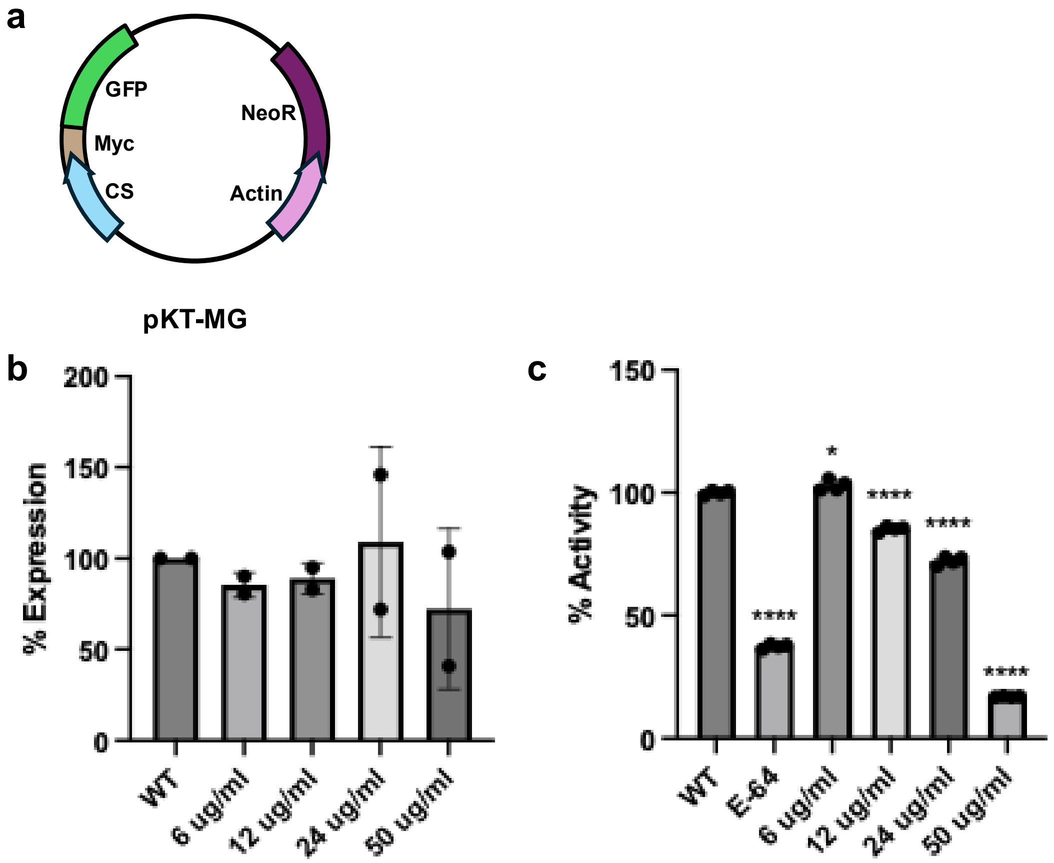

### Figure S3

#### Figure S3

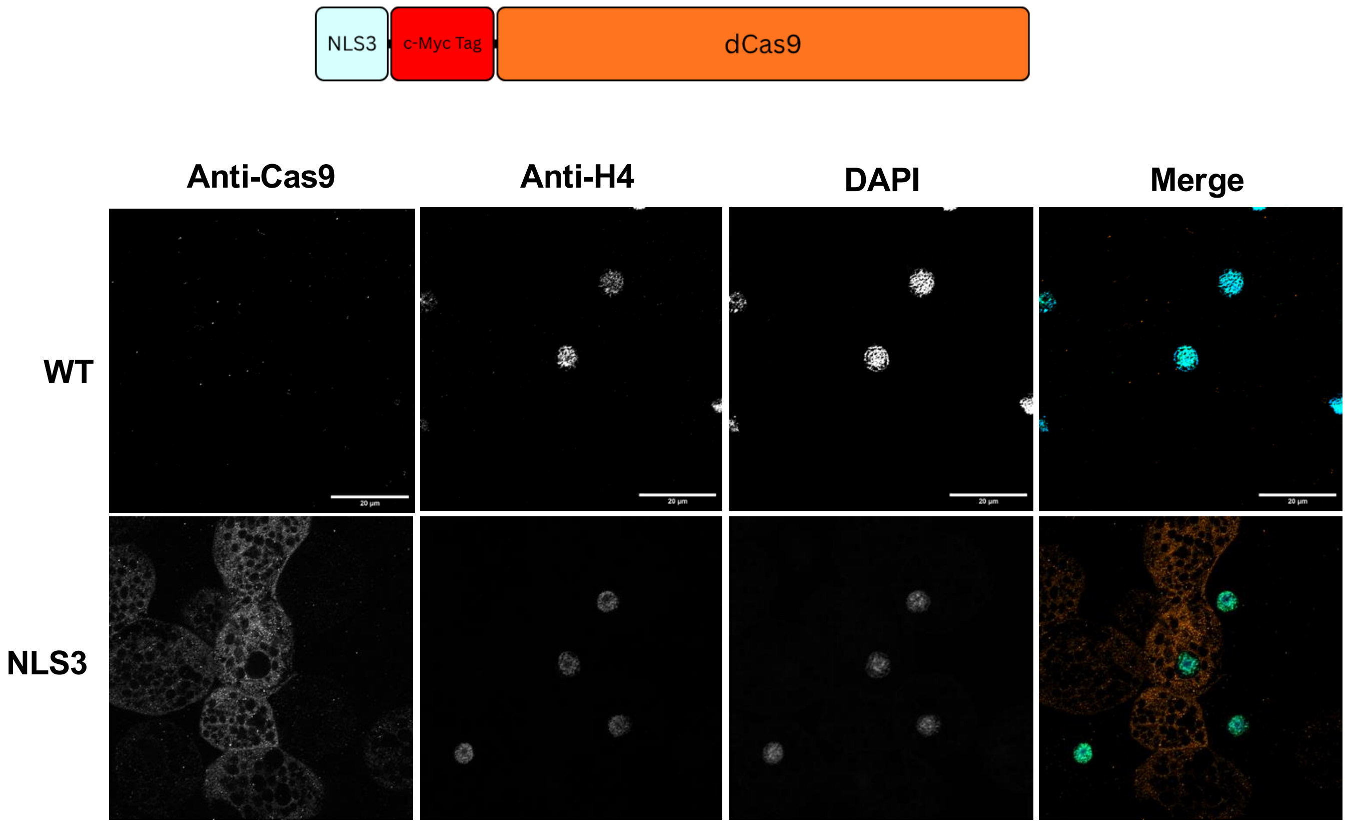

### Figure S4

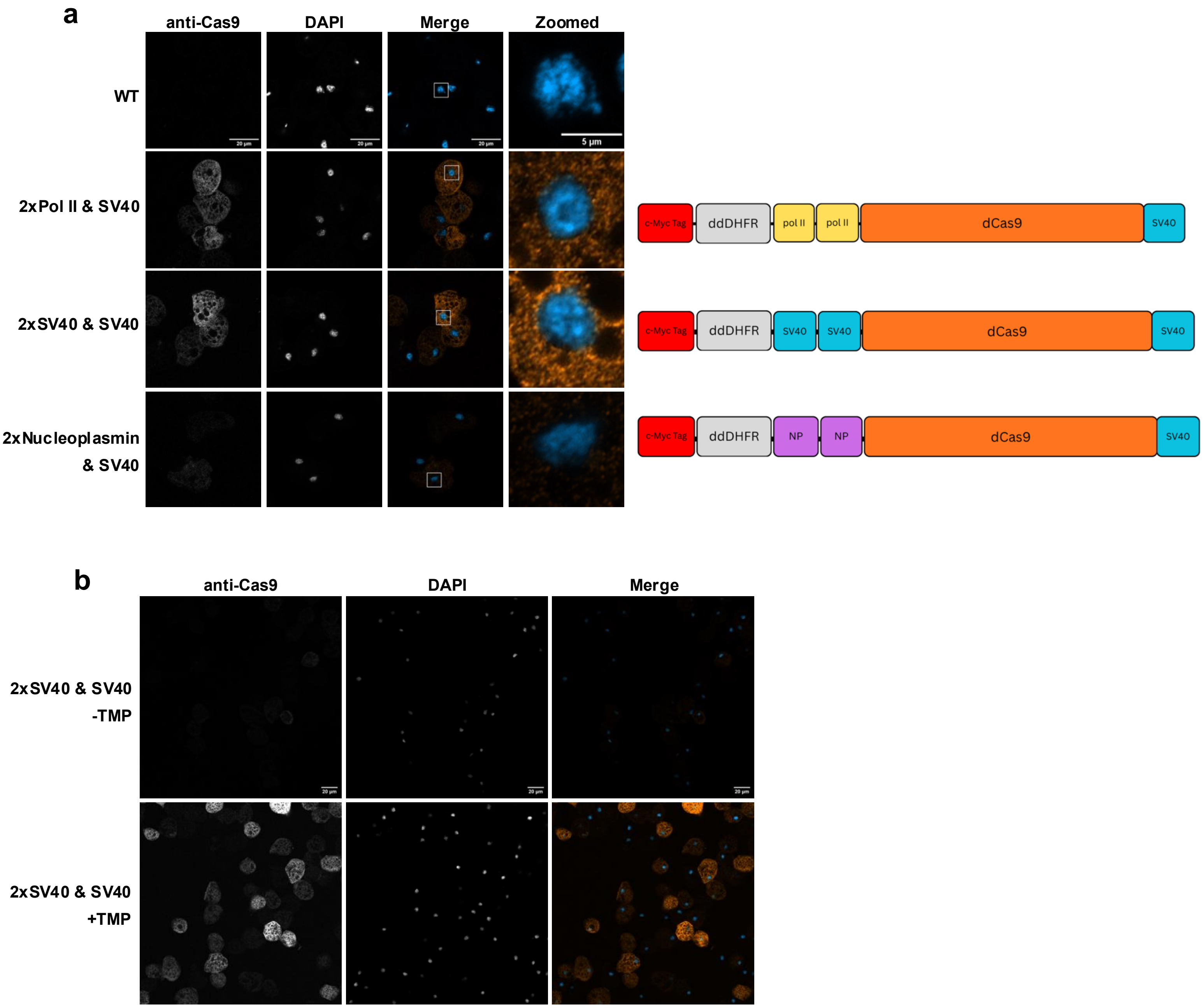

### Figure S5

#### Figure S5

a

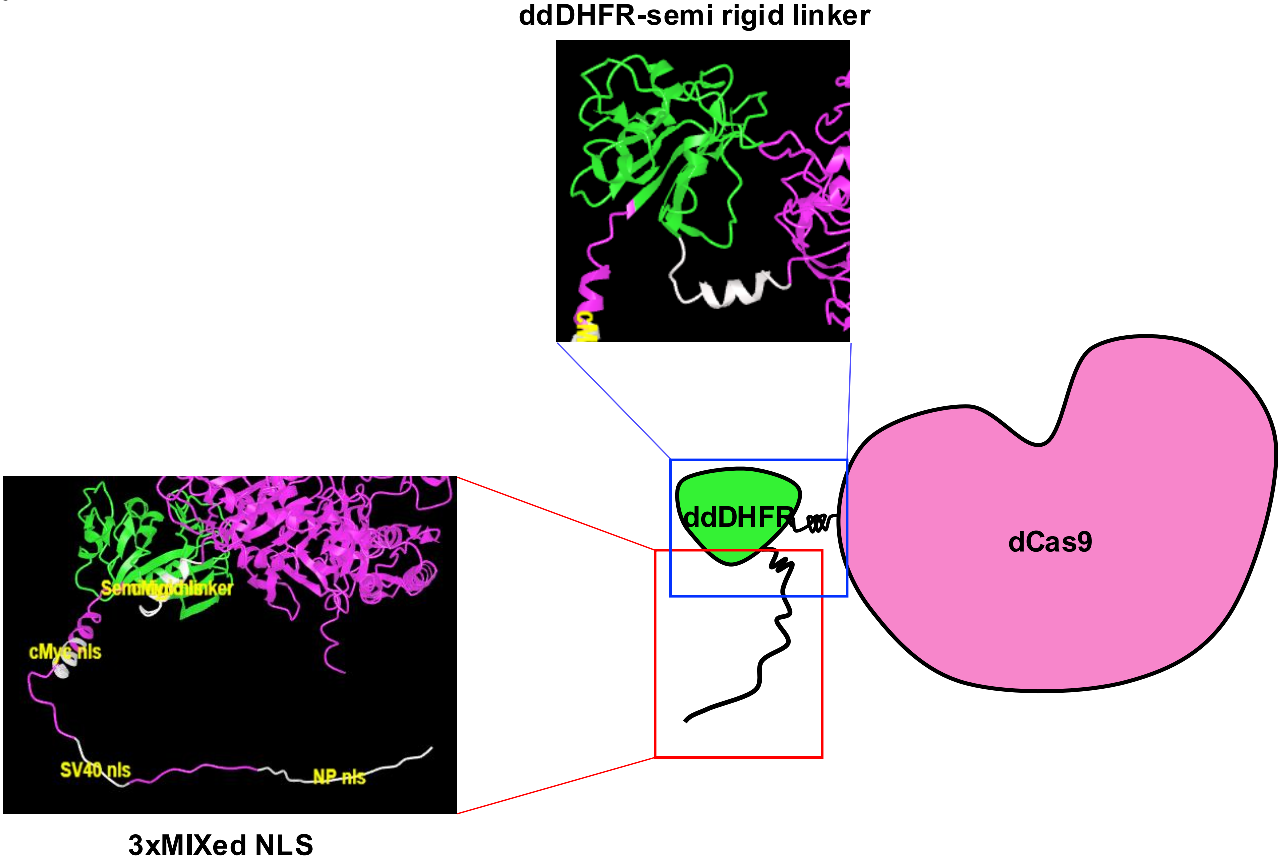


b

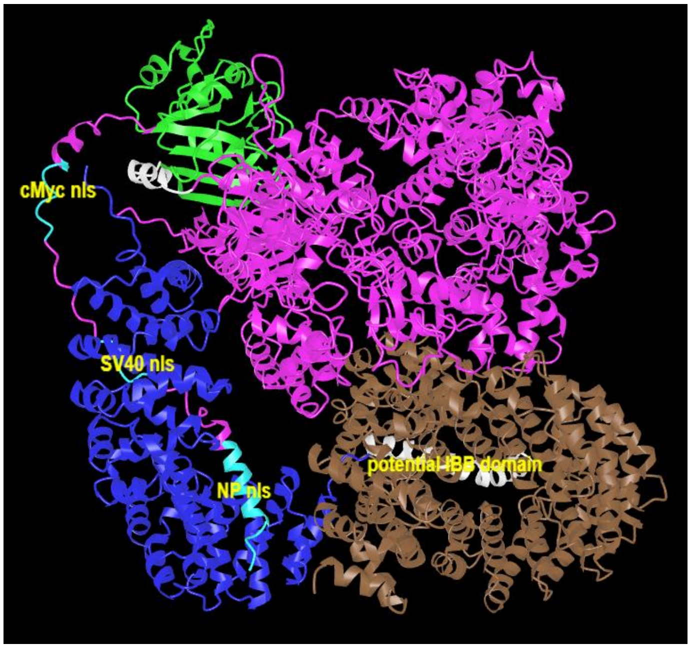


3xMIXed-ddDHFR-dCas9

Putative EhImportin alpha

Putative EhImportin beta

### Figure S6

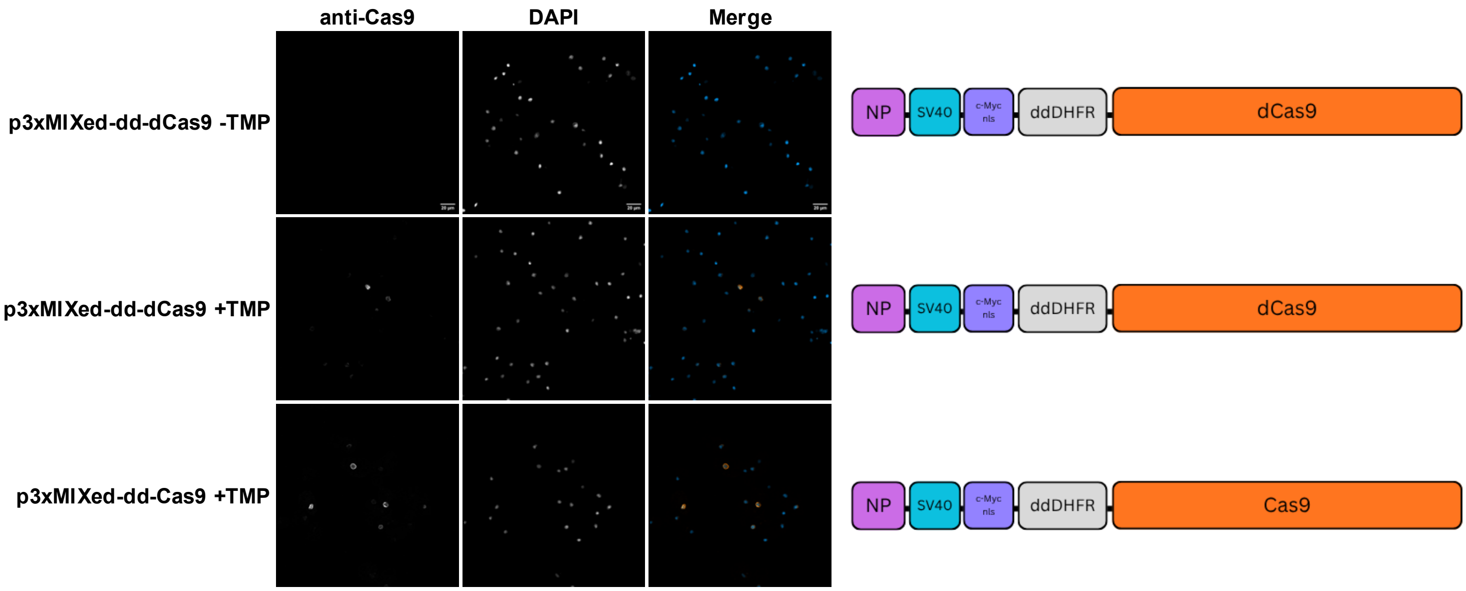

### Figure S7

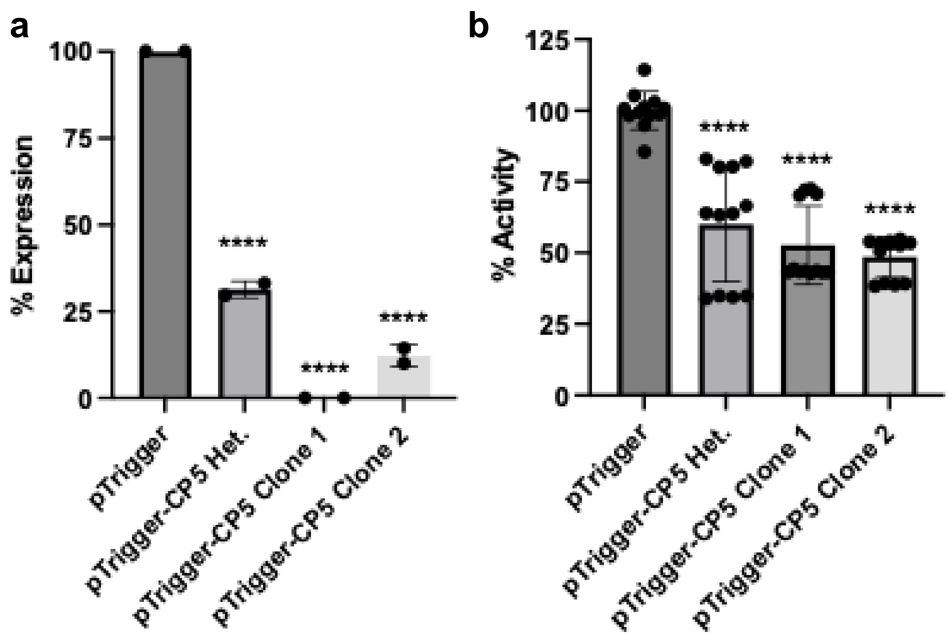

### Figure S8

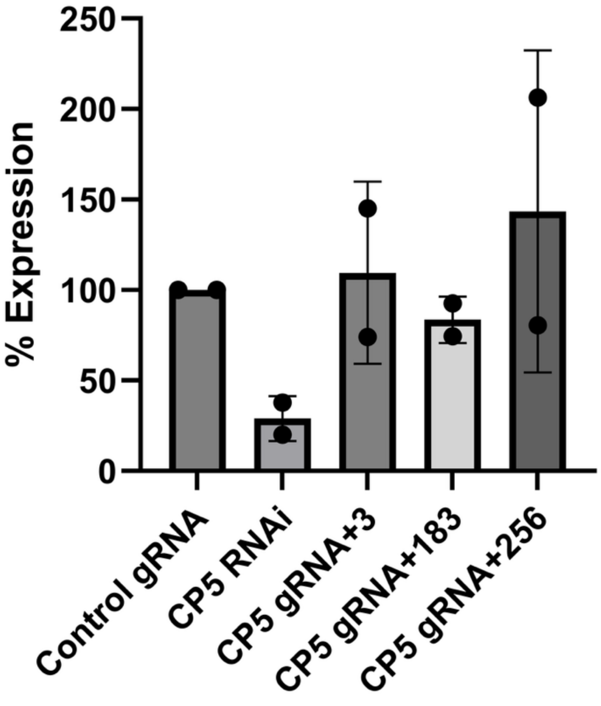

### Figure S9

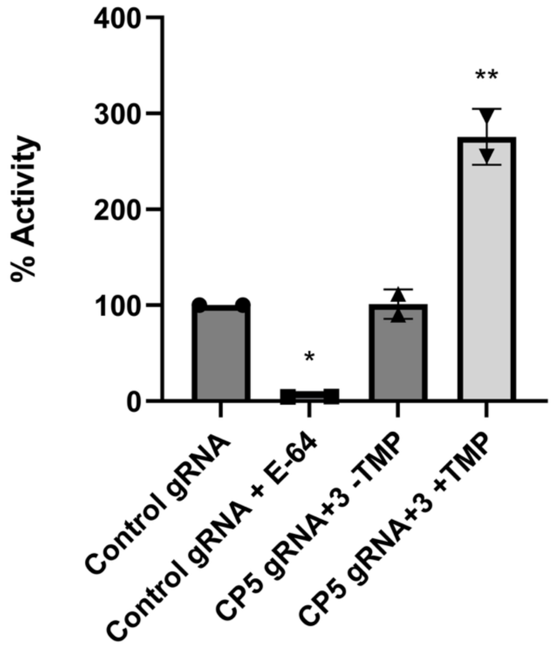
